# A pre-rRNA positive feedback loop drives malignant ribosome biogenesis

**DOI:** 10.64898/2026.07.03.736202

**Authors:** Emilie L. Alard, Martin Dodel, Shannon N. Russell, Zeinab Rekad, Maximilian Wiesbeck, Maia Cooper, Zaynah Patel, Muhammad S. Azman, Maria R. Conte, Stefan H. Stricker, Keaton I. Jones, Eric O’Neill, Faraz K. Mardakheh

## Abstract

Altered nucleoli are a well-established hallmark of cancer^1^, but how oncogenic signalling remodels the nucleolus remains poorly understood. Here we used an inducible mouse model of pancreatic ductal adenocarcinoma (PDAC)^2^ to generate spatially resolved proteomic and phosphoproteomic maps of the nucleolus upon RAS oncogene activation. We identify a phosphorylation programme initiated by translocation of the Casein Kinase 2 (CK2) holoenzyme to the nucleolus. This programme amplifies rRNA synthesis and malignant ribosome biogenesis by phosphorylating factors that control RNA polymerase I transcription and early ribosomal RNA (rRNA) processing. Preventing the nucleolar activity of CK2 inhibits oncogene-induced rRNA production, while constitutive nucleolar trapping of CK2 is sufficient to activate rRNA synthesis in the absence of RAS oncogene. Mechanistically, CK2 accumulation in the nucleolus is mediated by direct binding to the 3’ External Transcribed Sequence (3’ETS) of nascent precursor rRNA, creating an RNA-dependent self-amplifying feedback loop. Nucleolar CK2 accumulation is conserved across diverse human cancers, and its disruption synergises with inhibition of oncogenic RAS signalling to suppress anchorage-independent growth and tumourigenesis. Our study reveals 3’ETS as a CK2 signalling scaffold that amplifies oncogenic ribosome biogenesis, and defines a druggable nucleolar vulnerability that can be exploited by targeting this process.

## Introduction

The nucleolus is a dynamic membrane-less organelle within the nucleus that serves as the site of ribosome biogenesis^3^, one of the most resource-intensive anabolic processes in proliferating cells^4^. For more than a century, enlarged nucleoli have been recognised as a defining cytological feature of cancer^5^. Accordingly, nucleolar prominence is routinely used for both diagnosis as well as grading of diverse tumour types, with more prominent nucleolar staining frequently associated with aggressive disease and poor clinical outcome^6^. Enlarged nucleoli are also commonly observed in pre-cancerous lesions^7^, suggesting a close association with the driver mutations of cancer. Despite this long-standing association with tumourigenesis, the underlying molecular mechanisms that drive nucleolar enlargement, their relationship to the genetic drivers of malignancy, and the functional consequences of altered nucleolar architecture for tumorigenesis have remained unknown.

In human cells, nucleoli assemble after mitosis around nucleolar organiser regions (NORs), located on the short arms of the acrocentric chromosomes^8^. These regions contain hundreds of tandemly repeated ribosomal DNA (rDNA) loci, which are transcribed by RNA polymerase I (RNAPI) to generate the 47S precursor rRNA. This ∼13-kb transcript contains the 18S, 5.8S and 28S rRNA sequences, interspaced by internal and external transcribed spacers that are removed during the process of ribosome maturation. The fourth rRNA species, 5S rRNA, is transcribed independently by RNA polymerase III and imported into the nucleolus, where it becomes incorporated into the maturing large ribosomal subunit^9^. Ribosome biogenesis proceeds through a highly coordinated sequence of rRNA folding, modification, endonucleolytic cleavage, and exonucleolytic trimming events, coupled with the stepwise assembly of ribosomal proteins (RPs). This process is spatially organised across distinct nucleolar sub-compartments and requires hundreds of ribosome biogenesis factors (RBFs) and non-coding small nucleolar RNAs (snoRNAs)^9^. Importantly, internal and external transcribed spacer sequences have emerged as key organising elements that recruit and coordinate ribosome biogenesis factors^10–12^. However, the full extent of factors that bind these sequences, and how their assembly is dynamically coordinated during tumourigenesis, remains unclear.

Here, using an inducible mouse model of PDAC in which the expression of the driving Kras^G12D^ oncogene is controlled by doxycycline administration^2^, we show that oncogene induction triggers nucleolar fusion. This fusion depends on oncogenic RAS–MAPK signalling, and is accompanied by expansion of the fibrillar centre (FC) compartment of the nucleolus. Quantitative spatial proteomic and phosphoproteomic analyses revealed extensive remodelling of the nucleolar proteome following Kras^G12D^ activation, marked prominently by the nucleolar enrichment of CK2 holoenzyme, and widespread induction of CK2-dependent phosphorylation events. Notably, nucleolar accumulation of CK2 is conserved across diverse human cancers, indicating a broadly shared nucleolar programme underlying malignancy. Mechanistically, CK2 phosphorylates factors involved in RNA polymerase I transcription and early pre-rRNA processing within the FC compartment. This leads to amplification of rRNA synthesis, which in turn drives nucleolar fusion downstream of Kras^G12D^. Inhibiting CK2 activity suppresses oncogene-induced rRNA transcription and nucleolar fusion, whereas constitutive nucleolar sequestration of CK2 is sufficient to drive both processes independently of Kras^G12D^. We further show that CK2 accumulation within the nucleolus depends on its direct interaction with the 3′ETS sequence of the nascent pre-rRNA, establishing an RNA-dependent positive feedback loop that reinforces rRNA synthesis downstream of oncogenic signalling. Disruption of this pathway synergises with MAPK inhibition to suppress anchorage-independent growth and tumourigenesis of Kras^G12D^-driven PDAC cells, identifying a nucleolar therapeutic vulnerability in RAS-driven cancers. Taken together, we reveal a pre-rRNA-driven positive feedback mechanism that actively remodels the nucleolar structure and function to drive malignancy-associated nucleolar enlargement, ribosome biogenesis, and tumour growth, and demonstrate the possibility of therapeutically exploiting this process.

## Results

### Oncogenic RAS signalling induces nucleolar fusion

To investigate how malignancy reshapes the nucleolus, we chose PDAC as our disease model. Approximately 95% of PDAC cases are driven by activating mutations in KRAS, the most frequently mutated oncogene across all human cancers^13^. Analysis of matched normal, premalignant, and malignant human pancreatic tissue sections showed that nucleoli are frequently enlarged in both PDAC as well as pancreatic intraepithelial neoplasia (PanIN), the precursor lesions to PDAC that harbour KRAS mutations^14^, suggesting a causal link between the driver oncogenic RAS mutations and changes to the nucleolar morphology (Fig. 1a,b). To experimentally assess the impact of oncogenic RAS activation on the nucleolus, we used tumour cells isolated from the iKras PDAC mouse model^2^, in which the expression of Kras oncogene (Kras^G12D^) is controlled by doxycycline administration (Supplementary Data Fig. S1a). In the absence of Kras^G12D^, iKras cells behave akin to non-transformed cells, whereas Kras^G12D^ induction promotes transformation-associated phenotypes, including loss of contact inhibition (Supplementary Data Fig. S1b–d) and anchorage-independent growth (Supplementary Data Fig. S1e,f). Tumour formation *in vivo* is also strictly dependent on Kras^G12D^ expression (Supplementary Data Fig. S1g,h).

**Figure 1:**
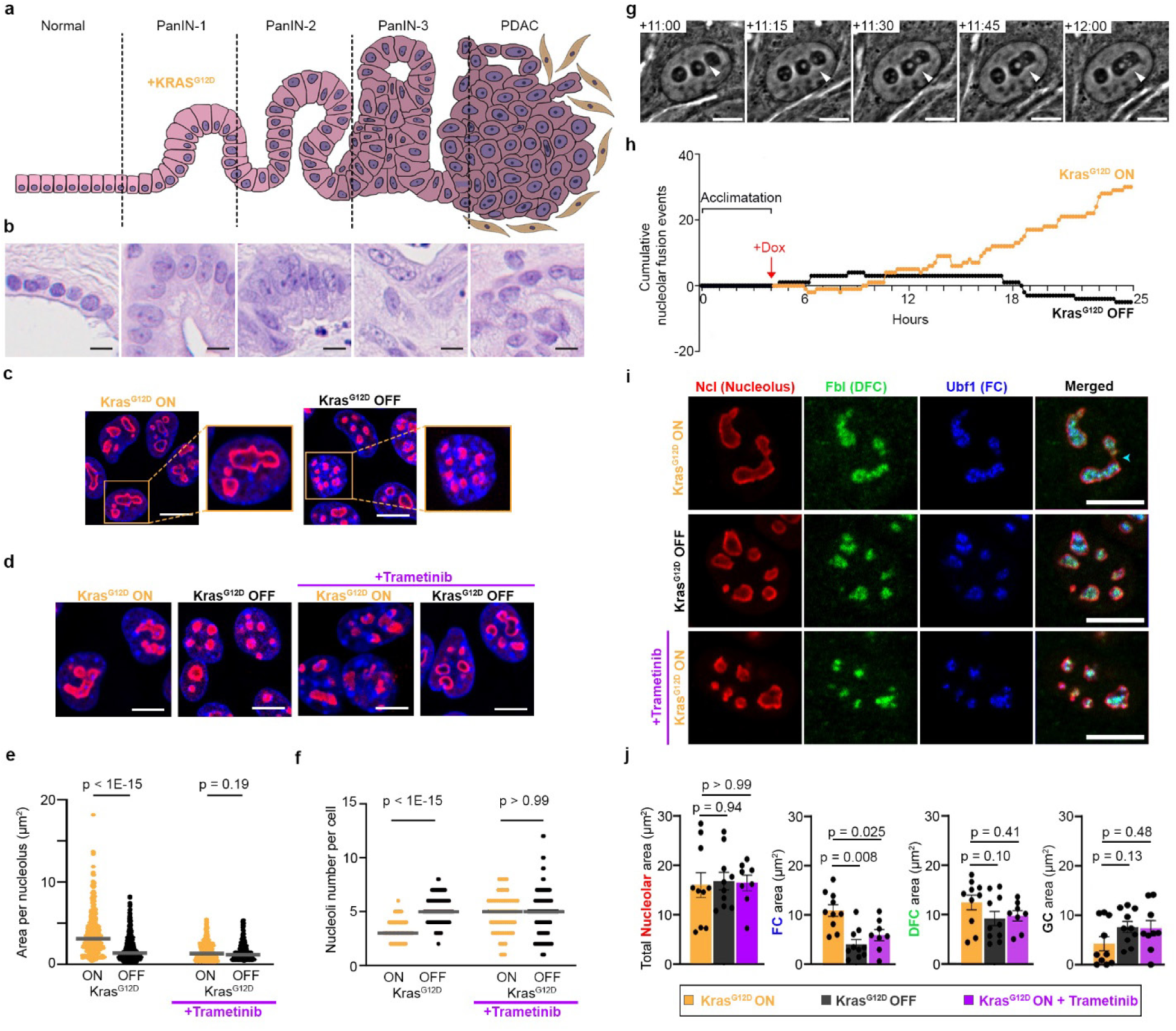
RAS–MAPK signalling promotes nucleolar fusion in PDAC. **(a)** Schematic diagram of PDAC progression from normal pancreatic epithelium, through premalignant PanIN-1, PanIN-2, and PanIN-3 pre-cancerous lesions, to invasive PDAC. Activating KRAS mutations, including KRAS^G12D^, are founding drivers of PanIN formation. **(b)** Representative H&E images selected from matched normal pancreas, PanIN, and PDAC human tissue samples, showing prominent nucleoli in low-, intermediate-, and high-grade PanINs, as well as in PDAC. A total of 20 matched normal and PDAC samples were stained, 6 of which also contained regions corresponding to various PanIN stages. Scale bar, 10 µm. **(c)** Representative immunofluorescence images of nucleoli in iKras PDAC cells treated with or without doxycycline for 24 h to induce Kras^G12D^ expression. Nucleoli were visualised by nucleolin (Ncl) immunostaining (red). Nuclei were stained with Hoechst (blue). Scale bar, 10 µm. **(d)** Representative immunofluorescence images of nucleoli in iKras PDAC cells treated with or without doxycycline for 24 h to induce Kras^G12D^ expression, with or without Trametinib (10 nM) co-treatment to inhibit MAPK signalling. Nucleoli were visualised by Ncl immunostaining (red). Nuclei were stained with Hoechst (blue). Scale bar, 10 µm. **(e)** Quantification of individual nucleolar area from the immunofluorescence images shown in (d). A total of n = 1,800 nucleoli pooled from three independent biological replicates were analysed. Significance was assessed by one-way ANOVA with Dunn’s multiple comparisons testing. **(f)** Quantification of nucleolar number per cell from the immunofluorescence images shown in (d). A total of n = 419 cells pooled from three independent biological replicates were analysed. Significance was assessed by one-way ANOVA with Dunn’s multiple comparisons testing. **(g)** Representative stills from phase-contrast live-cell imaging of iKras cells undergoing nucleolar fusion after doxycycline-induced Kras^G12D^ expression. Cells were acclimatised in the imaging chamber for 4 h before doxycycline addition, followed by 21 h of imaging. The indicated fusion event occurs 11–12 h after doxycycline addition. White arrow indicates the fusion event. Scale bar, 5 µm. **(h)** Quantification of cumulative nucleolar fusion (+1) and fission (−1) events from phase-contrast live-cell imaging of iKras cells described in (g). A total of n = 40 cells pooled from three independent biological replicates were analysed. **(i)** Representative immunofluorescence images of nucleolar sub-compartments in iKras PDAC cells treated with or without doxycycline for 24 h, with or without Trametinib co-treatment. Whole nucleoli were visualised by Ncl immunostaining, the DFC by fibrillarin (Fbl) immunostaining, and the FC by upstream binding factor 1 (Ubf1) immunostaining. Scale bar, 10 µm. **(j)** Quantification of total nucleolar area, along with FC, DFC, and GC areas per cell, from the immunofluorescence images shown in (i). GC area was estimated by subtracting DFC area from total nucleolar area. A total of n = 38 cells pooled from three independent biological replicates were analysed. Significance was assessed by two-way ANOVA with Šídák’s multiple comparisons testing.

Immunofluorescence analysis of iKras cells revealed that induction of Kras^G12D^ expression triggered nucleolar enlargement (Fig. 1c). This effect was dependent on RAS–MAPK signalling, the primary effector pathway downstream of Kras^G12D^ in these cells^15^, as it was abolished by trametinib, a potent small-molecule inhibitor of MAPK signalling^16^ (Fig. 1d). Notably, Kras^G12D^ withdrawal or MAPK inhibition reduced the individual nucleolar size but increased the number of nucleoli per cell (Fig. 1e,f), suggesting that RAS–MAPK signalling is promoting the coalescence of smaller nucleoli into larger structures. Consistent with this model, live-cell imaging revealed that Kras^G12D^ induction triggers widespread nucleolar fusion (Fig. 1g,h; Video S1). This process began approximately 6 h after doxycycline administration, the time-point coinciding with the effective Kras^G12D^ expression and MAPK activation in these cells^17^, and continued progressively thereafter (Fig. 1h). Immunofluorescence analysis of nucleolar sub-compartments revealed that RAS-induced nucleolar fusion did not significantly alter the overall sizes of the granular compartment (GC) or dense fibrillar compartment (DFC), but significantly increased the size of the fibrillar centres (FC) (Fig. 1i,j). As FCs are closely associated with rDNA transcription, their enlargement suggests that in addition to nucleolar fusion, oncogenic RAS–MAPK signalling induces a reorganisation of the nucleolar architecture, potentially linked to altered rDNA transcription.

### Oncogenic RAS signalling localises CK2 to the nucleolus

Whilst prior studies have carried out proteomics characterisation of nucleolar composition^18^, it is unclear how the nucleolar content is dynamically regulated in response to signalling. To determine how oncogenic RAS signalling remodels the nucleolar composition, we devised a quantitative spatial proteomics approach (Fig. 2a). Nucleoli were isolated via sucrose cushion fractionation^19^ from iKras cells cultured with or without Kras^G12D^ expression, or with Kras^G12D^ expression together with Trametinib co-treatment to inhibit RAS–MAPK signalling (Supplementary Data Fig. S2a). In parallel, matched whole-cell lysates (WCLs) were prepared from the same conditions. All samples were then trypsin digested and analysed by tandem mass tag (TMT)-based quantitative proteomics and phosphoproteomics (Fig. 2a).

**Figure 2:**
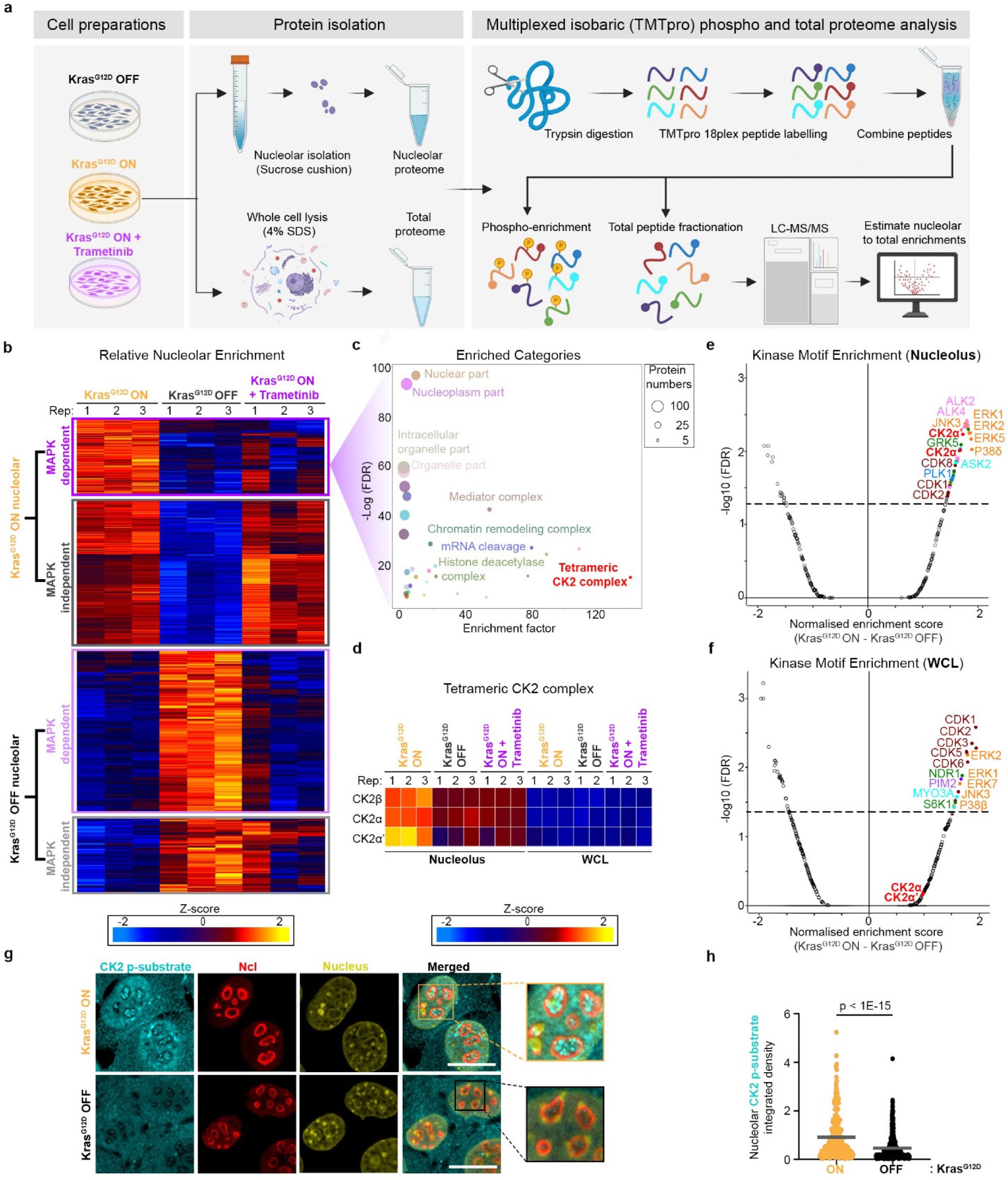
Nucleolar proteomics reveals RAS–MAPK-dependent localisation of CK2 signalling. **(a)** Workflow for quantitative proteomic and phosphoproteomic profiling of the nucleolus. Three independent biological replicates were analysed per condition. **(b)** Heatmap of proteins whose nucleolar levels relative to the WCL was significantly increased or decreased upon 24h of Kras^G12D^ induction in iKras cells, in a MAPK-dependent or-independent manner, as judged by sensitivity to co-treatment with Trametinib (10 nM). Significant changes were identified using one-way ANOVA with permutation-based FDR of < 0.05 and S0 = 0.1, followed by post hoc testing with FDR of < 0.05 (Dataset S4). **(c)** Category enrichment analysis of proteins recruited to the nucleolus upon Kras^G12D^ induction in a MAPK-dependent manner from (b). Enriched categories were identified using Fisher’s exact test with a Benjamini–Hochberg FDR cut-off of < 0.001, enrichment factor > 2, and intersection size > 2 (Dataset S5). Each circle represents an enriched category from Gene Ontology Cellular Component (GOCC) or *Dorner et al.* ribosome biogenesis annotations^9^; circle size indicates the number of intersecting proteins. **(d)** Heatmap showing relative levels of CK2α, CK2α, and CK2β, in the nucleolus and WCL samples of different treatment conditions. **(e)** Volcano plot of normalised enrichment scores for serine/threonine kinase motifs^23^ in the nucleolus upon Kras^G12D^ induction (Dataset S8). Dashed line: adjusted p-value < 0.05. **(f)** Volcano plot of normalised enrichment scores for serine/threonine kinase motifs^23^ in the WCL upon Kras^G12D^ induction (Dataset S9). Dashed line: adjusted p-value < 0.05. **(g)** Representative immunofluorescence images of CK2 phosphorylated substrates (cyan) in iKras cells treated with or without doxycycline for 24 h to induce Kras^G12D^ expression. Nucleoli were visualised by Ncl immunostaining (red), while Nuclei were stained by Hoechst (yellow). Insets show higher-magnification views of selected nucleoli. Scale bar, 10 µm. **(h)** Quantification of nucleolar CK2-phosphorylated substrate levels from the immunofluorescence images shown in (g). A total of n = 690 nucleoli pooled from three independent biological replicates were analysed. Significance was assessed using a two-tailed, unpaired, non-parametric Mann–Whitney test.

Principal component analysis (PCA) of the total proteome showed clear separation between nucleolar and WCL fractions, as well as a largely expected clustering of biological replicates within each treatment group (Supplementary Data Fig. S2b,c). Comparison of nucleolar fractions with WCLs showed strong enrichment of known nucleolar protein markers (Supplementary Data Fig. S2d; Dataset S1), as well as whole categories of nucleolar annotated proteins, including several families of RBFs (Supplementary Data Fig. S2e; Dataset S2). These data confirmed successful capture of the nucleolar proteome. We next assessed how Kras^G12D^ signalling alters the nucleolar proteome. ANOVA coupled with post hoc analysis revealed that of 5,153 quantified proteins, 2,130 were enriched in the nucleolus in at least one treatment condition (Dataset S3). Although the nucleolar enrichment of most proteins was unaffected by Kras^G12D^ expression, 291 proteins were significantly regulated by Kras^G12D^ in a MAPK-dependent manner, whereas 599 proteins were regulated independently of MAPK signalling (Supplementary Data Fig. S2f). Post hoc analysis of Kras^G12D^-responsive nucleolar proteins further identified four clusters whose nucleolar enrichment either increased or decreased following Kras^G12D^ induction, in either a MAPK-dependent or MAPK-independent manner (Fig. 2b; Dataset S4).

Category enrichment analysis identified the tetrameric CK2 complex among the most strongly enriched protein categories that localised to the nucleolus in a RAS–MAPK-dependent manner (Fig. 2c; Dataset S5). Several ribosome biogenesis-related protein categories were also recruited to the nucleolus following Kras^G12D^ induction, but in a MAPK-independent manner (Supplementary Data Fig. S2g; Dataset S5). In contrast, protein categories associated with chromatin compaction and remodelling were depleted from the nucleolus upon Kras^G12D^ induction (Supplementary Data Fig. S2h,i; Dataset S5), suggesting that oncogenic RAS signalling may promote a reorganisation of nucleolar chromatin.

CK2 is a highly conserved, constitutively active kinase composed of two catalytic subunits, CK2α and/or CK2α′, and two regulatory CK2β subunits. It regulates diverse cellular processes, including signal transduction, DNA repair, and cell division^20^. CK2 also interacts with and phosphorylates multiple components of the RNAPI pre-initiation complex^21^. Our proteomics analysis revealed significant RAS–MAPK-dependent enrichment of all CK2 subunits in the nucleolus (Fig. 2d). Moreover, our previous work had showed that Kras^G12D^ induction promotes phosphorylation of CK2α at T360 and S362, modifications reported to increase CK2 kinase activity by 3 to 4-fold^22^. We therefore hypothesised that MAPK-dependent recruitment of CK2 to the nucleolus, together with its enhanced activity downstream of MAPK, would specifically increase the phosphorylation of nucleolar substrates. To test this, we next analysed the nucleolar phosphoproteome dynamics in response RAS–MAPK signalling. As observed for the total proteome, PCA of the phosphoproteome showed clear separation between nucleolar and WCL fractions, as well as clustering of replicates within their treatment groups (Supplementary Data Fig. S2j). Comparison of nucleolar phosphoproteomes with matched WCLs showed enrichment of phosphorylated nucleolar markers (Supplementary Data Fig. S2k; Dataset S6), as well as phosphorylated proteins belonging to nucleolar functional classes, including several families of RBFs (Supplementary Data Fig. S2l; Dataset S7), thus confirming the successful capture of the nucleolar phosphoproteome.

We then used the Kinase Library motif enrichment analysis^23^ to infer kinase activities that were altered by Kras^G12D^ induction. CK2 and MAPK (a.k.a. ERK) were among the top predicted kinases activated downstream of Kras^G12D^ within the nucleolus (Fig. 2e; Dataset S8). In contrast, CK2 activity was not significantly increased in WCLs (Fig. 2f; Dataset S9), suggesting that RAS–MAPK-dependent CK2 recruitment enhances CK2 activity specifically within the nucleolar compartment. Consistent with this, immunofluorescence analysis of iKras cells with an antibody that recognises CK2-phosphorylated substrates revealed a significant accumulation of CK2-phosphorylated proteins within the nucleolus following Kras^G12D^ induction (Fig. 2g,h). Together, these findings reveal that oncogenic RAS–MAPK signalling recruits CK2 to the nucleolus, where it establishes a compartment-specific nucleolar phosphorylation programme.

### CK2 localisation triggers nucleolar fusion

Next, we set out to validate the localisation of CK2 to the nucleolus and investigate its functional significance. Immunofluorescence analysis of iKras cells with a CK2β-specific antibody confirmed the RAS–MAPK-dependent nucleolar enrichment of CK2β (Fig. 3a,b; Supplementary Data Fig. S3a,b). Importantly, immunohistochemistry (IHC) analysis of human pancreatic tissues revealed that CK2β was also enriched in nucleoli of human PanIN and PDAC cells, indicating that this localisation is conserved in humans (Fig. 3c,d). Furthermore, analysis of CK2α IHC images from the Human Protein Atlas^24^ revealed cancer-specific nucleolar localisation of CK2α across several tumour types (Supplementary Data Fig. S3c,d), suggesting that nucleolar enrichment of CK2 may be a broader feature of malignancy.

**Figure 3:**
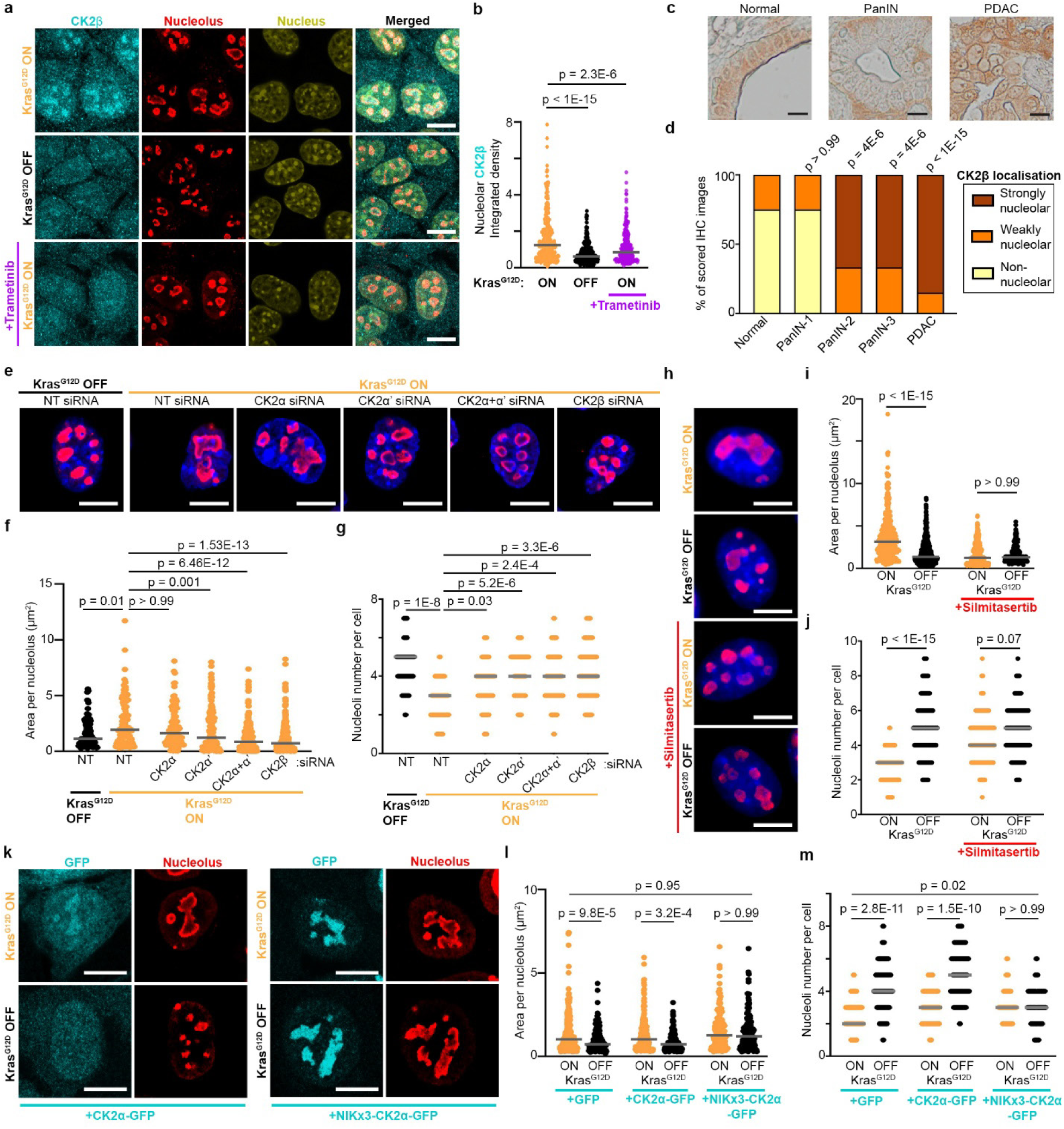
Kras^G12D^-dependent localisation of CK2 to the nucleolus drives nucleolar fusion. **(a)** Representative immunofluorescence images of CK2β (cyan) in iKras cells treated with or without doxycycline for 24 h to induce Kras^G12D^ expression, or with doxycycline plus Trametinib (10 nM) co-treatment to inhibit MAPK signalling. Nucleoli were visualised by Ncl immunostaining (red). Nuclei were stained with Hoechst (yellow). Scale bar, 10 µm. **(b)** Quantification of nucleolar CK2β levels from the immunofluorescence images shown in (a). A total of n = 932 nucleoli pooled from three independent biological replicates were analysed. Significance was assessed by one-way ANOVA with Dunn’s multiple comparisons testing. **(c)** Representative IHC images of matched normal, PanIN, and PDAC human tissue samples stained for CK2β (brown). A total of 20 matched normal and PDAC samples were analysed, 6 of which also contained regions corresponding to distinct PanIN stages. Scale bar, 10 µm. **(d)** Quantification of nucleolar CK2β staining patterns in tissue images described in (c). Images were scored as either no nucleolar, weakly nucleolar, or strongly nucleolar. Multiple regions were imaged per sample. Significance was assessed by Fisher’s exact test. Normal, n = 80 images; PanIN-1, n = 16 images; PanIN-2, n = 6 images; PanIN-3, n = 6 images; PDAC, n = 60 images. **(e)** Representative immunofluorescence images of nucleoli in iKras cells transfected with indicated siRNAs for 72 h, with or without doxycycline for 24 h to induce Kras^G12D^ expression. Nucleoli were visualised by Ncl immunostaining (red). Nuclei were stained with Hoechst (blue). Scale bar, 10 µm. **(f)** Quantification of individual nucleolar area from the immunofluorescence images shown in (e). A total of n = 968 nucleoli pooled from three independent biological replicates were analysed. Significance was assessed by one-way ANOVA with Dunn’s multiple comparisons testing. **(g)** Quantification of nucleolar number per cell from the immunofluorescence images shown in (e). A total of n = 262 cells pooled from three independent biological replicates were analysed. Significance was assessed by one-way ANOVA with Dunn’s multiple comparisons testing. **(h)** Representative immunofluorescence images of nucleoli in iKras cells treated with or without doxycycline for 24 h to induce Kras^G12D^ expression, with or without Silmitasertib (10 µM) co-treatment to inhibit CK2 kinase activity. Nucleoli were visualised by Ncl immunostaining (red). Nuclei were stained with Hoechst (blue). Scale bar, 10 µm. **(i)** Quantification of individual nucleolar area from the immunofluorescence images shown in (h). A total of n = 1775 nucleoli pooled from three independent biological replicates were analysed. Significance was assessed by one-way ANOVA with Dunn’s multiple comparisons testing. **(j)** Quantification of nucleolar number per cell from the immunofluorescence images shown in (h). A total of n = 403 cells pooled from three independent biological replicates were analysed. Significance was assessed by one-way ANOVA with Dunn’s multiple comparisons testing. **(k)** Representative immunofluorescence images of iKras cells ectopically expressing GFP-tagged (cyan) wild-type CK2α, or NIKx3-fused CK2α, together with untagged CK2β. Cells were treated with or without doxycycline for 24 h to induce Kras^G12D^ expression. Nucleoli were visualised by Ncl immunostaining (red). Nuclei were stained with Hoechst (yellow). Scale bar, 10 µm. **(l)** Quantification of individual nucleolar area from the immunofluorescence images shown in (k). A total of n = 1041 nucleoli pooled from three independent biological replicates were analysed. Significance was assessed by one-way ANOVA with Dunn’s multiple comparisons testing. **(m)** Quantification of nucleolar number per cell from the immunofluorescence images shown in (k). A total of n = 361 cells pooled from three independent biological replicates were analysed. Significance was assessed by one-way ANOVA with Dunn’s multiple comparisons testing.

Next, we assessed the impact of CK2 activity in regulating nucleolar morphology. RNAi-mediated depletion of CK2 subunits, either individually or in combination, was performed using subunit-specific siRNAs (Supplementary Data Fig. S3e). Depletion of CK2β, or combined depletion of the catalytic CK2α and CK2α′ subunits, abolished Kras^G12D^-induced nucleolar fusion, with CK2α′ depletion alone also significantly reducing fusion but to a lesser extent (Fig. 3e–g). Given that CK2α′ showed a stronger nucleolar enrichment than CK2α (Fig. 2d), these results suggested that the catalytic activity of CK2 within the nucleolus could be required for RAS-induced nucleolar fusion. Consistent with this notion, treatment with Silmitasertib, a selective small-molecule inhibitor of CK2^25^, completely abolished nucleolar fusion downstream of Kras^G12D^ (Fig. 3h–j). To assess if localising CK2 to the nucleolus is sufficient to trigger nucleolar fusion, we next generated expression constructs for GFP-tagged CK2α, with or without fusion to three tandem nucleolar targeting sequences from the NF-κB-inducing kinase (NIKx3)^26^. While GFP-tagged wild-type CK2α accumulated in the nucleolus in a Kras^G12D^-dependent manner similar to the endogenous protein, NIKx3-tagged CK2α was constitutively nucleolar, irrespective of Kras^G12D^ (Supplementary Data Fig. S3f,g). Crucially, constitutive retention of CK2α in the nucleolus was sufficient to induce nucleolar fusion irrespective of Kras^G12D^ (Fig. 3k–m), suggesting that CK2 localisation is the main driver of nucleolar fusion downstream of oncogenic RAS signalling.

### Localised CK2 activity promotes nucleolar fusion by stimulating rRNA synthesis

Next, we set out to determine the role of nucleolar CK2 downstream of oncogenic RAS signalling. To identify the nucleolar phosphorylation targets of CK2 in an unbiased manner, we performed another quantitative spatial proteomics profiling in iKras cells cultured with or without Kras^G12D^ expression, or with Kras^G12D^ expression together with short-term Silmitasertib treatment. We reasoned that a 1-hour Silmitasertib treatment would preferentially capture the more immediate CK2-dependent phosphorylation events, thereby enriching for the direct targets of CK2 kinase activity. As before, nucleolar fractions and matched whole-cell lysates (WCLs) were prepared from each condition, trypsin-digested, and analysed by TMT-based quantitative proteomics and phosphoproteomics.

PCA of the total proteome showed clear separation between nucleolar and WCL fractions, consistent with successful fractionation (Supplementary Data Fig. S4a). A similar separation was observed in the phosphoproteome PCA (Supplementary Data Fig. S4b). ANOVA followed by post hoc analysis of the total proteome identified only 63 proteins whose nucleolar enrichments were dependent on CK2 activity (Supplementary Data Fig. S4c; Dataset S10), consistent with the short duration of Silmitasertib treatment. Notably, CK2 subunits themselves were among the proteins whose nucleolar enrichment was significantly reduced by short-term Silmitasertib treatment (Supplementary Data Fig. S4d), suggesting that kinase activity of CK2 is critical for its own accumulation in the nucleolus.

We next analysed the changes to the nucleolar phosphoproteome. Of 7,825 identified phosphorylation sites, ANOVA followed by post hoc analysis identified 2,098 sites to be specifically enriched in the nucleolus (Dataset S11). Among these, 368 phosphorylation sites were enriched in a Kras^G12D^ and CK2-dependent manner, whilst 880 were Kras^G12D^-dependent but CK2-independent (Fig. 4a; Dataset S12). Thus, CK2 accounts for approximately one-third of Kras^G12D^-dependent nucleolar phosphorylation events. As expected, the CK2 motif was the most significantly enriched kinase motif among Silmitasertib-sensitive phosphorylation sites, supporting the likely identification of direct CK2 targets (Fig. 4b; Dataset S13). Category enrichment analysis showed that Kras^G12D^ and CK2-dependent nucleolar phosphorylation targets were enriched in proteins involved in rDNA transcription and early ribosomal processing (Fig. 4c; Dataset S14). By contrast, Kras^G12D^-dependent but CK2-independent phosphorylation targets were enriched for various diverse protein groups involved in both early and late steps of ribosome biogenesis (Supplementary Data Fig. S4e; Dataset S14).

**Figure 4:**
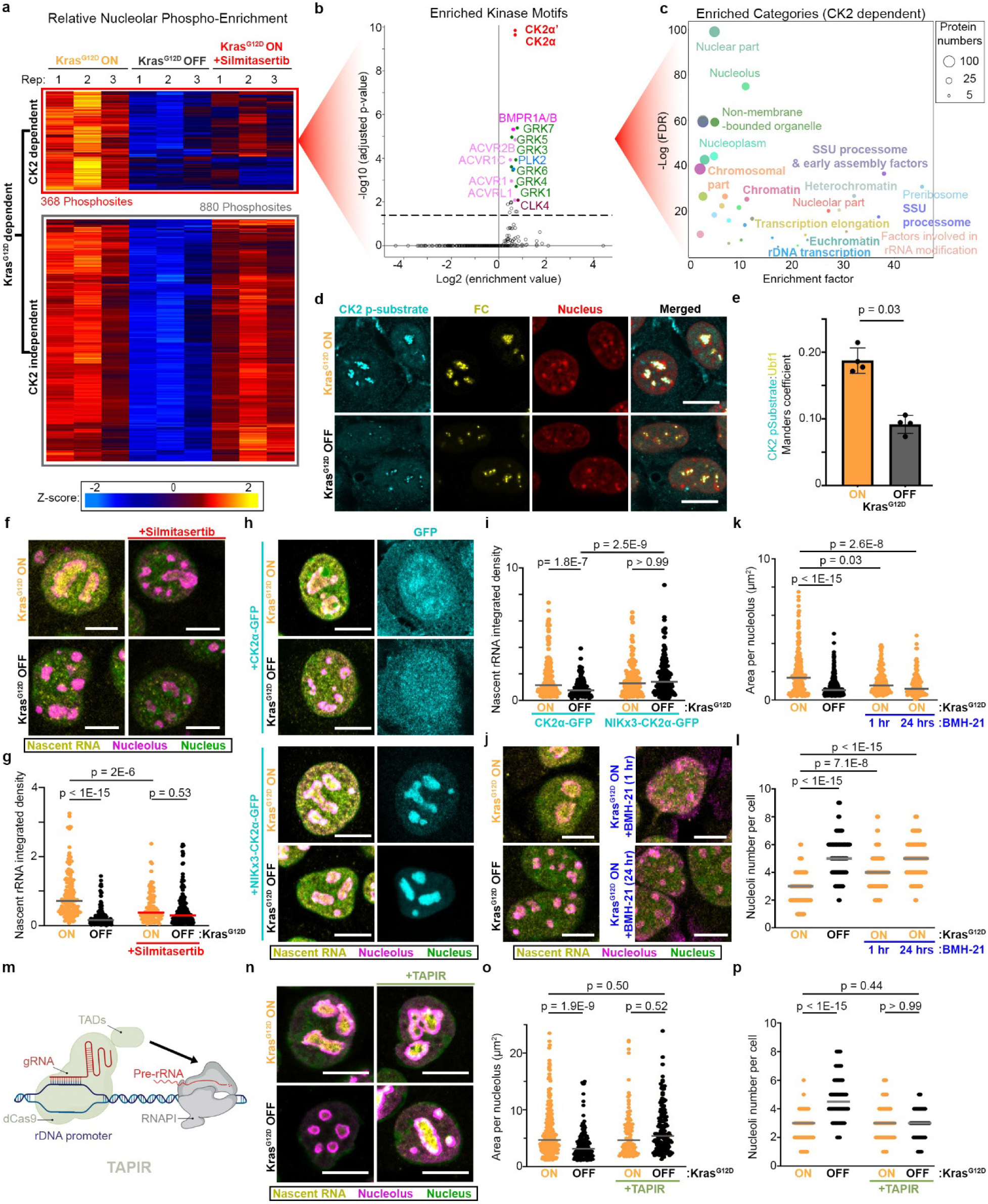
Nucleolar CK2 activity triggers nucleolar fusion by promoting rRNA synthesis. **(a)** Heatmap of phosphorylation sites whose nucleolar levels relative to the WCL was significantly increased upon 24 h of Kras^G12D^ induction in iKras cells, in a CK2-dependent or CK2-independent manner, as judged by sensitivity to 1 h of Silmitasertib (10 µM) treatment. Significant changes were identified using one-way ANOVA with permutation-based FDR of < 0.05 and S0 = 0.1, followed by post hoc testing with FDR of < 0.05 (Dataset S12). **(b)** Fisher enrichment analysis of serine/threonine kinase motifs^23^ amongst the Kras^G12D^ and CK2-dependent nucleolar phosphorylation sites from (a) (Dataset S13). (Dashed line: adjusted p-value < 0.05). **(c)** Category enrichment analysis of Kras^G12D^ and CK2-dependent nucleolar phosphorylation targets from (a). Enriched categories were identified using Fisher’s exact test with a Benjamini–Hochberg FDR cut-off of < 0.02, enrichment factor > 2, and intersection size > 2 (Dataset S14). Each circle represents an enriched category from Gene Ontology Cellular Component (GOCC) or *Dorner et al.* ribosome biogenesis annotations^9^; circle size indicates the number of intersecting proteins. **(d)** Representative immunofluorescence images of CK2 phosphorylated substrates (cyan) colocalisation with the FC sub-compartment of the nucleolus in iKras cells treated with or without doxycycline for 24 h to induce Kras^G12D^ expression. The FC sub-compartment was visualised by Ubf1 immunostaining (yellow). Nuclei were stained by Hoechst (red). Scale bar, 10 µm. **(e)** Manders coefficients for the fraction of CK2 phosphorylated substrates colocalising with the FC sub-compartment from immunofluorescence images shown in (d). Average coefficients from n = 4 fields of view per condition, each containing 7–11 cells, were quantified. Significance was calculated using a two-sided two-sample t-test. **(f)** Representative nascent RNA immunofluorescence images of iKras cells treated with or without doxycycline for 24 h to induce Kras^G12D^ expression, with or without Silmitasertib (10 µM) co-treatment to inhibit CK2 activity. Cells were pulse-labelled with FUrd (2 mM) for 30 min to label nascent RNA, before immunostaining for FUrd (yellow). Nucleoli were visualised by Ncl immunostaining (red). Nuclei were stained with Hoechst (green). Scale bar, 10 µm. **(g)** Quantification of nucleolar FUrd levels from the immunofluorescence images shown in (f). A total of n = 739 nucleoli pooled from three independent biological replicates were analysed. Significance was assessed by one-way ANOVA with Dunn’s multiple comparisons testing. **(h)** Representative nascent RNA immunofluorescence images of iKras cells ectopically expressing GFP-tagged wild-type CK2α or NIKx3-fused CK2α (cyan), together with untagged CK2β. Cells were treated with or without doxycycline for 24 h to induce Kras^G12D^ expression, pulse-labelled with FUrd (2 mM) for 30 min to label nascent RNA, before immunostaining for FUrd (yellow). Nucleoli were visualised by Ncl immunostaining (red). Nuclei were stained with Hoechst (green). Scale bar, 10 µm. **(i)** Quantification of nucleolar FUrd levels from the immunofluorescence images shown in (h). A total of n = 639 nucleoli pooled from three independent biological replicates were analysed. Significance was assessed by one-way ANOVA with Dunn’s multiple comparisons testing. **(j)** Representative nascent RNA immunofluorescence images of iKras cells treated with or without doxycycline for 24 h to induce Kras^G12D^ expression, with or without BMH-21 (0.5 µM) co-treatment for indicated times to inhibit rRNA synthesis. Cells were pulse-labelled with FUrd (2 mM) for 30 min to label nascent RNA, before immunostaining for FUrd (yellow). Nucleoli were visualised by Ncl immunostaining (red). Nuclei were stained with Hoechst (green). Scale bar, 10 µm. **(k)** Quantification of individual nucleolar area from the immunofluorescence images shown in (j). A total of n = 907 nucleoli pooled from three independent biological replicates were analysed. Significance was assessed by one-way ANOVA with Dunn’s multiple comparisons testing. **(l)** Quantification of nucleolar number per cell from the immunofluorescence images shown in (j). A total of n = 445 cells pooled from three independent biological replicates were analysed. Significance was assessed by one-way ANOVA with Dunn’s multiple comparisons testing. **(m)** Schematic diagram of TAPIR^31^. Specific gRNAs are used to target a dCas9 protein fused to three transcriptional activation domains (TADs) to the rDNA promoter regions, leading to RNAPI transcription activation. **(n)** Representative nascent RNA immunofluorescence images of iKras cells with or without ectopic expression of TAPIR. Indicated cells were treated with or without doxycycline for 24 h to induce Kras^G12D^ expression, pulse-labelled with FUrd (2 mM) for 30 min to label nascent RNA, before immunostaining for FUrd (yellow). Nucleoli were visualised by Ncl immunostaining (red). Nuclei were stained with Hoechst (green). Scale bar, 10 µm. **(o)** Quantification of individual nucleolar area from the immunofluorescence images shown in (n). A total of n = 701 nucleoli pooled from three independent biological replicates were analysed. Significance was assessed by one-way ANOVA with Dunn’s multiple comparisons testing. **(p)** Quantification of nucleolar number per cell from the immunofluorescence images shown in (n). A total of n = 429 cells pooled from three independent biological replicates were analysed. Significance was assessed by one-way ANOVA with Dunn’s multiple comparisons testing.

Amongst the CK2-independent targets, we identified two phosphorylation sites on Ubf1, the master transcription factor for RNAPI, consistent with previous reports that MAPK-dependent phosphorylation of Ubf1 activates rDNA transcription^27^ (Supplementary Data Fig. S4f). In contrast, CK2-dependent targets included other components of the RNAPI transcription machinery, including Rrn3, Tcof1, and Ttf1^9^, as well as Ncl, which we previously showed to enhance rRNA synthesis upon phosphorylation by CK2^17^ (Supplementary Data Fig. S4g). Several regulators of nucleolar chromatin organisation, including Baz1b^28^, Myc^29^, and Rrp8^30^, were also identified amongst the CK2-dependent phosphorylation targets (Supplementary Data Fig. S4g). Together, these data suggest that nucleolar CK2 activity specifically regulates rDNA organisation and RNAPI transcription. Consistent with this model, immunofluorescence analysis using the antibody against CK2-phosphorylated substrates revealed a strong colocalisation of nucleolar CK2 phosphorylation targets with the FC sub-compartment, with increased overlap following Kras^G12D^ induction (Fig. 4d,e).

Next, we examined the effect of CK2 kinase activity on rRNA synthesis downstream of Kras^G12D^. Nascent rRNA was visualised by pulse-labelling cells with the uridine analogue 5-fluorouridine (FUrd), followed by antibody-based detection of nucleolar FUrd-incorporated RNA^32^. As reported previously, induction of Kras^G12D^ expression strongly enhanced rRNA synthesis^17^; however, this effect was abolished by Silmitasertib treatment, indicating that CK2 kinase activity is required for Kras^G12D^-induced stimulation of RNAPI transcription (Fig. 4f,g). To determine whether nucleolar localisation of CK2 is sufficient to promote rRNA synthesis, we ectopically expressed GFP-tagged wild-type CK2α or the constitutively nucleolar NIKx3-fused variant in iKras cells, with or without subsequent Kras^G12D^ induction. Constitutive retention of CK2α in the nucleolus activated rRNA synthesis irrespective of Kras^G12D^ status (Fig. 4h,i), indicating that localisation of CK2 to the nucleolus is sufficient to enhance rRNA synthesis.

Since rDNA transcription emerged as a major downstream consequence of nucleolar CK2 activity, we next tested whether enhanced rRNA synthesis was behind the Kras^G12D^-induced nucleolar fusions. Inhibition of rRNA synthesis with BMH-21, a small-molecule RNAPI inhibitor^33^, for as little as 1 hour significantly reduced Kras^G12D^-induced nucleolar fusion, with 24 hours of treatment causing a further reduction (Fig. 4j–l; Supplementary Data Fig. S4h). Similar results were obtained after 24 hours of treatment with CX-5461, an independent RNAPI inhibitor^34^ (Supplementary Data Fig. S4i–l). As a complementary approach, we activated rRNA synthesis independently of any upstream signalling by using TAPIR (Targeted Activation of Protein Translation), our recently developed CRISPR activation-based tool to ectopically increase rRNA synthesis^31^ (Fig. 4m). TAPIR expression led to constitutive activation of rRNA synthesis irrespective of Kras^G12D^ status (Fig. 4n; Supplementary Data Fig. S4m). Crucially, this was sufficient to induce nucleolar fusion in the absence of Kras^G12D^ (Fig. 4n–p). Together, these results reveal that oncogenic RAS–MAPK signalling recruits CK2 to the nucleolus, where it specifically phosphorylates several key components of rDNA transcription machinery, resulting in an increase in rRNA synthesis. This increase in rRNA production in turn promotes nucleolar fusion, leading to emergence of prominent nucleoli.

### 3’ETS rRNA recruits CK2 to the nucleolus

We next set out to determine how oncogenic RAS–MAPK signalling targets CK2 to the nucleolus. Since CK2 nucleolar enrichment was reduced by acute inhibition of its kinase activity (Supplementary Data Fig. S4d), we reasoned that a CK2-dependent nucleolar process might be reinforcing CK2 localisation. As our data identified rDNA transcription as a major downstream output of nucleolar CK2 activity, we asked whether RNAPI activity was required for CK2 recruitment to the nucleolus. Inhibition of RNAPI transcription by BMH-21 or CX-5461, for as short as one hour, abolished Kras^G12D^-induced nucleolar enrichment of CK2, while ectopic activation of rRNA synthesis using TAPIR was sufficient to drive CK2 accumulation in the nucleolus independently of Kras^G12D^ (Fig. 5a–d). Nucleolar CK2 activity showed the same dependence on RNAPI transcription, with BMH-21 and CX-5461 preventing the Kras^G12D^-induced nucleolar accumulation of CK2-phosphorylated substrates, whilst TAPIR increasing their accumulation regardless of Kras^G12D^ expression (Supplementary Data Fig. S5a–d). These results indicate that active rDNA transcription is required for CK2 localisation and activity within the nucleolus.

**Figure 5:**
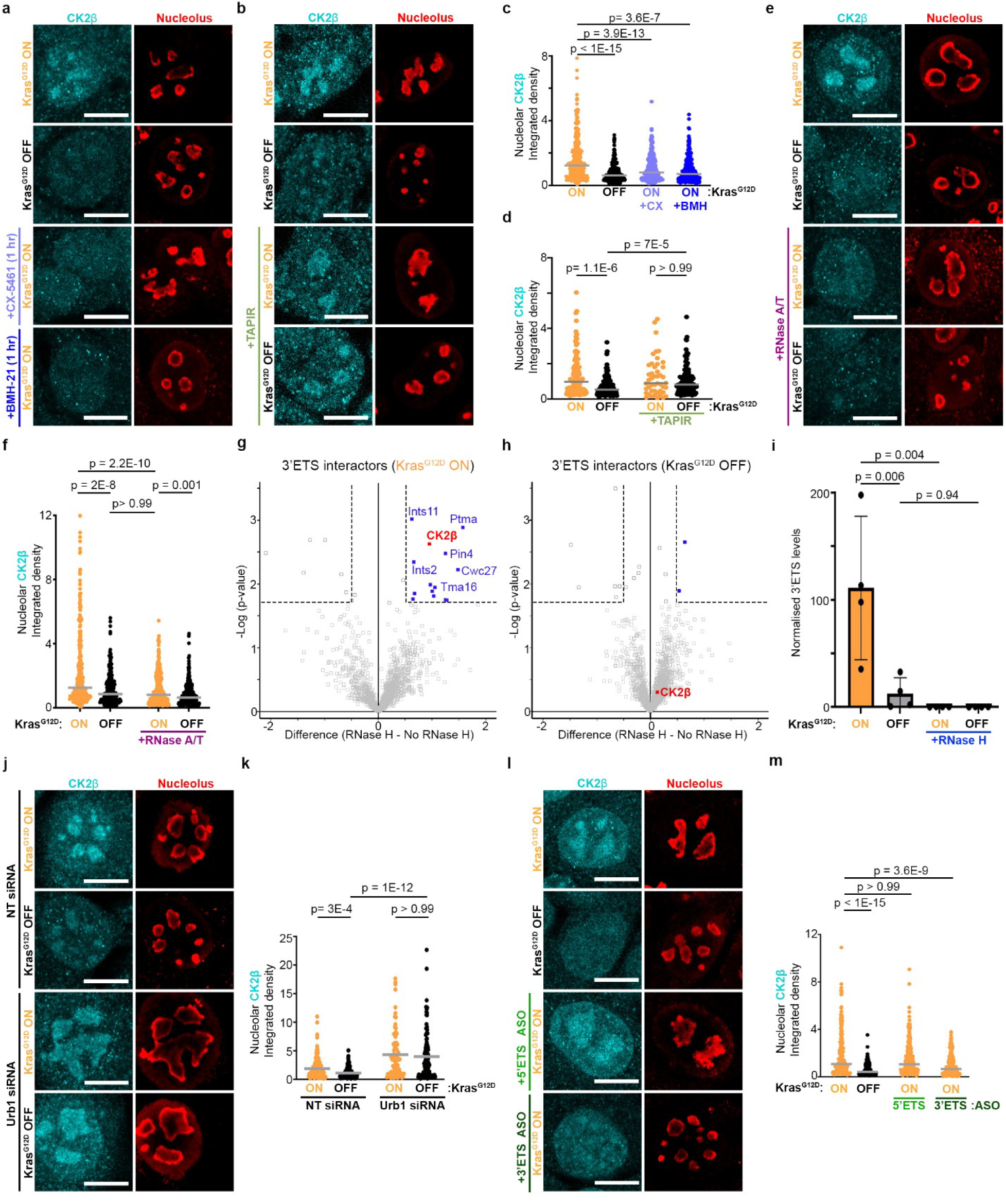
3’ETS rRNA binds to CK2 and enriches it in the nucleolus downstream of Kras^G12D^. **(a)** Representative immunofluorescence images of CK2β (cyan) in iKras cells treated with or without doxycycline for 24 h to induce Kras^G12D^ expression, or with doxycycline plus 1 h co-treatment with CX-5461 (100 nM) or BMH-21 (500 nM) to inhibit RNAPI. Nucleoli were visualised by Ncl immunostaining (red). Scale bar, 10 µm. **(b)** Representative immunofluorescence images of CK2β (cyan) in iKras cells treated with or without doxycycline for 24 h to induce Kras^G12D^ expression, with or without TAPIR expression to ectopically activate RNAPI. Nucleoli were visualised by Ncl immunostaining (red). Scale bar, 10 µm. **(c)** Quantification of nucleolar CK2β levels from the immunofluorescence images shown in (a). A total of n = 1200 nucleoli pooled from three independent biological replicates were analysed. Significance was assessed by one-way ANOVA with Dunn’s multiple comparisons testing. **(d)** Quantification of nucleolar CK2β levels from the immunofluorescence images shown in (b). A total of n = 415 nucleoli pooled from three independent biological replicates were analysed. Significance was assessed by one-way ANOVA with Dunn’s multiple comparisons testing. **(e)** Representative immunofluorescence images of CK2β (cyan) in iKras cells treated with or without doxycycline for 24 h to induce Kras^G12D^ expression, with or without 1 unit of RNase A+T1 treatment for 1 h post-fixation to remove RNA-based localisations. Nucleoli were visualised by Ncl immunostaining (red). Scale bar, 10 µm. **(f)** Quantification of nucleolar CK2β levels from the immunofluorescence images shown in (e). A total of n = 1138 nucleoli pooled from three independent biological replicates were analysed. Significance was assessed by one-way ANOVA with Dunn’s multiple comparisons testing. **(g)** Volcano plot comparison of 3’ETS-digested (n = 4) versus undigested (n = 3) TREX samples from iKras cells treated with doxycycline for 24 h to induce Kras^G12D^ expression. Significance was assessed using a two-sided, two-sample t-test. Hits were defined based on an absolute difference score > 0.5 and p < 0.02 (Dataset S15). **(h)** Volcano plot comparison of 3’ETS-digested (n = 4) versus undigested control (n = 3) TREX samples of iKras cells cultured without doxycycline. Significance was assessed using a two-sided, two-sample t-test. Hits were defined based on an absolute difference score > 0.5 and p < 0.02 (Dataset S15). **(i)** RT-qPCR analysis of 3′ETS RNA levels in RNase H-digested and undigested TREX samples from (g) and (h). A 10% fraction of each TREX sample (n = 4) was used for RNA extraction and RT-qPCR analysis. 3′ETS RNA levels were normalised to the average of β-Actin and 18S RNAs as house-keeping controls. Significance was assessed by one-way ANOVA with Holm-Šídák’s multiple comparisons testing. **(j)** Representative immunofluorescence images of CK2β (cyan) in iKras cells transfected with Urb1-targeting or non-targeting control siRNAs, with or without doxycycline for 24 h to induce Kras^G12D^ expression. Nucleoli were visualised by Ncl immunostaining (red). Scale bar, 10 µm. **(k)** Quantification of nucleolar CK2β levels from the immunofluorescence images shown in (j). A total of n = 504 nucleoli pooled from three independent biological replicates were analysed. Significance was assessed by one-way ANOVA with Dunn’s multiple comparisons testing. **(l)** Representative immunofluorescence images of CK2β (cyan) in iKras cells mock transfected or transfected with ASOs targeting the 5’ETS or 3’ETS, with or without doxycycline for 24 h to induce Kras^G12D^ expression. Nucleoli were visualised by Ncl immunostaining (red). Scale bar, 10 µm. **(m)** Quantification of nucleolar CK2β levels from the immunofluorescence images shown in (l). A total of n = 985 nucleoli pooled from three independent biological replicates were analysed. Significance was assessed by one-way ANOVA with Dunn’s multiple comparisons testing.

Since Kras^G12D^ induction increases the accumulation of nascent pre-rRNA in the nucleolus, we next investigated whether pre-rRNA itself contributes to CK2 recruitment. Post-fixation RNase treatment of iKras cells significantly reduced the Kras^G12D^-dependent nucleolar accumulation of CK2, indicating that RNA is actively required to maintain CK2 in the nucleolus (Fig. 5e,f). Importantly, we had recently mapped the direct region-by-region protein interactome of pre-rRNA in RAS-mutant human HCT116 cells by using Targeted RNase H-mediated extraction of crosslinked RBPs (TREX)^35^. TREX combines UV crosslinking and organic phase separation, with sequence-specific RNase H-mediated RNA digestion to release and identify proteins that directly associate with defined RNA regions in living cells^35^. Mining this previously generated human pre-rRNA interactome map revealed CK2α amongst the specific interactors of the 3′ETS region (Supplementary Data Fig. S5e,f), suggesting that the 3′ETS could be acting as the RNA element that retains CK2 in the nucleolus through direct binding.

To validate this interaction in the iKras system, we next performed a quantitative TREX analysis of the 3′ETS in iKras cells with or without Kras^G12D^ expression. PCA demonstrated a clear separation between Kras^G12D^-expressing and non-expressing samples (Supplementary Data Fig. S5g). In Kras^G12D^-expressing cells, 14 specific 3′ETS interactors were identified, which included CK2β as well as ribosome biogenesis factors such as Tma16 and Pin4 (Fig. 5g; Dataset S15). Importantly, most 3′ETS interactions, including CK2β, were lost upon Kras^G12D^ removal (Fig. 5h; Dataset S15). This loss was likely due to a ∼10-fold reduction in 3′ETS abundance in cells without Kras^G12D^ expression (Fig. 5i). This dramatic difference in 3’ETS levels with or without Kras^G12D^ expression is consistent with early studies suggesting this spacer region to be amongst the earliest to be processed following transcription termination^36^, making its abundance likely to closely scale with 47S rRNA synthesis rates.

Unlike CK2β, CK2α was not consistently detected across all RNase H-treated iKras TREX replicates, so its interaction with 3′ETS could not be confidently validated in these cells. Nevertheless, direct binding of CK2β to the mouse 3′ETS, together with our previous identification of CK2α as a 3′ETS-specific interactor in human cells, suggests that the CK2 holoenzyme likely engages with mammalian 3′ETS-containing pre-rRNAs. Together, these results support a model in which oncogenic RAS–MAPK signalling increases nascent pre-rRNA production, generating 3′ETS-containing transcripts that directly bind CK2.

We next tested whether 3′ETS was functionally required for CK2 recruitment to the nucleolus. For this purpose, we used two orthogonal approaches. First, we increased the abundance of 3′ETS-containing transcripts by depleting Urb1, a recently identified factor required for 3′ETS cleavage and degradation^37^. As expected, Urb1 depletion markedly increased the levels of 3′ETS (Supplementary Data Fig. S5h). Importantly, this increase was sufficient to drive a strong nucleolar accumulation of CK2, irrespective of Kras^G12D^ expression (Fig. 5j,k). Urb1 depletion also triggered nucleolar fusions, as judged by a significant increase in the nucleolar area combined with a significant decrease in nucleolar number (Supplementary Data Fig. S5i,j). Second, we disrupted 3′ETS interactions in living cells using antisense oligonucleotides (ASOs)^38^. ASOs targeting the full length of 3′ETS, but not 5′ETS, significantly impaired CK2 accumulation in the nucleolus downstream of Kras^G12D^ (Fig. 5l,m), along with blocking Kras^G12D^-induced nucleolar fusions (Supplementary Data Fig. S5k,l). Together, these results indicate that the 3′ETS region of pre-rRNA acts as a recruitment scaffold for CK2, enabling its nucleolar accumulation, phosphorylation of downstream targets, and induction of nucleolar fusion downstream of RAS oncogene.

### CK2 binding to 3’ETS creates a positive feedback loop that can be therapeutically exploited

The dependence of CK2 localisation on 3′ETS suggests a positive feedback mechanism linking oncogenic RAS signalling to hyperactive ribosome biogenesis. In this model, an initial RAS–MAPK signal stimulates rRNA synthesis by phosphorylating key activators of rDNA transcription, such as Ubf1, thereby increasing production of nascent 3′ETS-containing pre-rRNA. 3’ETS subsequently binds to and promotes CK2 accumulation in the nucleolus, where it phosphorylates additional components of rDNA chromatin and RNAPI transcription machinery, triggering further amplification of rRNA synthesis rates downstream of oncogenic RAS signalling (Fig. 6a).

**Figure 6:**
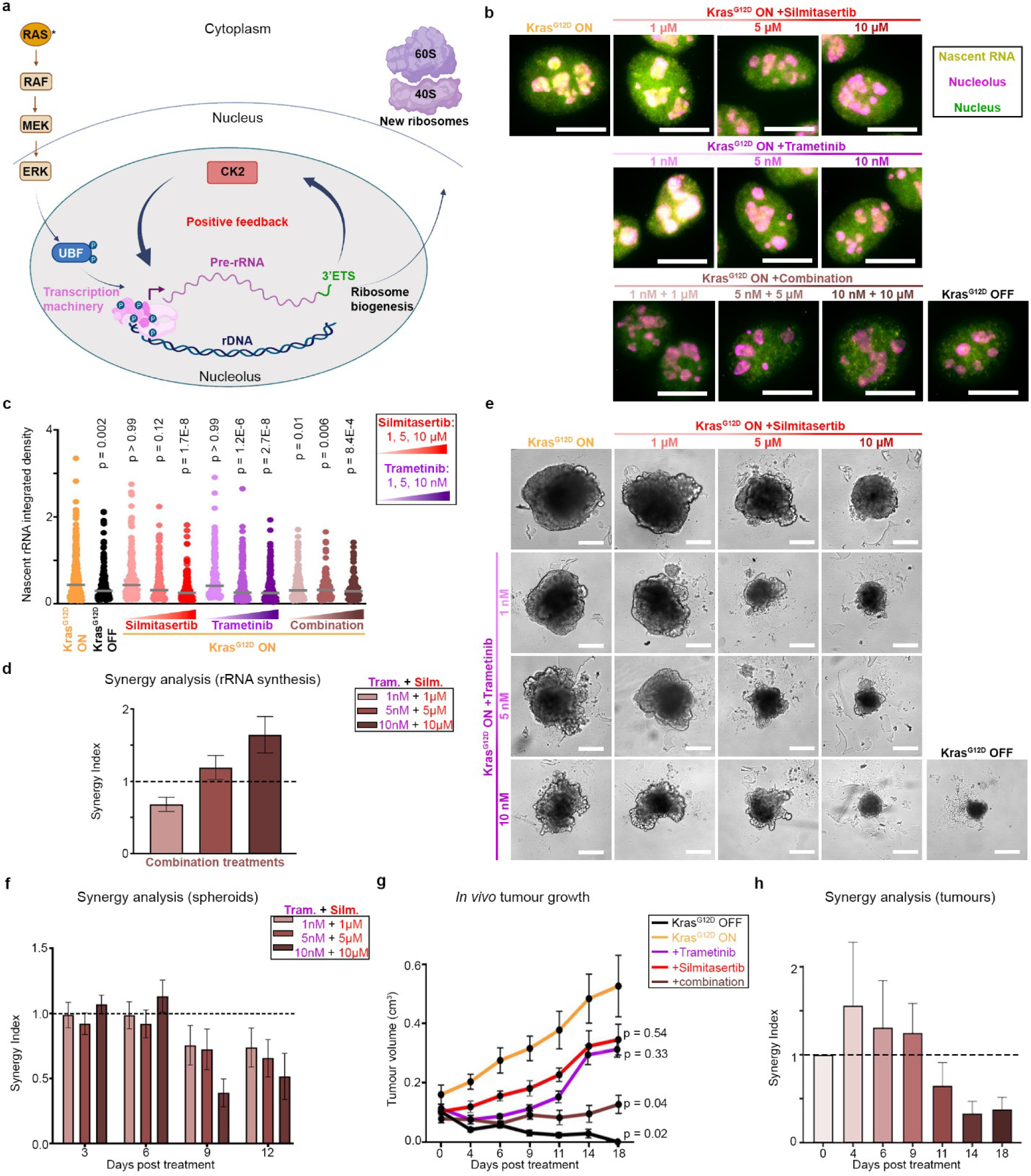
3’ETS-mediated nucleolar recruitment of CK2 creates a therapeutically targetable positive feedback loop that amplifies rRNA synthesis downstream of oncogenic RAS signalling. **(a)** Schematic diagram of the proposed positive feedback model in which oncogenic RAS signalling amplifies ribosome biogenesis through 3′ ETS-mediated recruitment of CK2 to the nucleolus. **(b)** Representative nascent RNA immunofluorescence images of iKras cells treated with or without doxycycline for 24 h to induce Kras^G12D^ expression, or with doxycycline plus co-treatment for 1 h with the indicated doses of Trametinib and Silmitasertib, either alone or in combination. Cells were pulse-labelled with FUrd (2 mM) for 30 min to label nascent RNA, before immunostaining for FUrd (yellow). Nucleoli were visualised by Ncl immunostaining (red). Nuclei were stained with Hoechst (green). Scale bar, 10 µm. **(c)** Quantification of nucleolar FUrd levels from the immunofluorescence images shown in (b). A total of n = 2,003 nucleoli pooled from three independent biological replicates were analysed. Significance was assessed by one-way ANOVA with Dunn’s multiple comparisons testing. **(d)** Synergy index (SI) for inhibition of nascent RNA synthesis by combined Trametinib and Silmitasertib treatment, calculated from the quantifications shown in (c). Error bars were calculated by error propagation using S.E.M. values. SI < 1 indicates synergy. **(e)** Representative images of iKras 3D spheroids cultured with or without doxycycline, or with doxycycline plus co-treatment with the indicated doses of Trametinib and Silmitasertib, either alone or in combination. Spheroids were established for 3 days in the presence of doxycycline, then subjected to the indicated treatments for 3, 6, 9, or 12 days. Selected images show spheroids after 12 days of treatment. Scale bar, 200 µm. **(f)** Synergy index (SI) for inhibition of 3D spheroid growth by combined Trametinib and Silmitasertib treatment, calculated from the quantifications shown in (Supplementary Data Fig. S6c). Error bars were calculated by error propagation using S.E.M. values. SI < 1 indicates synergy. **(g)** Tumour growth curves of subcutaneous iKras xenografts in nude mice, subjected to the indicated treatment conditions. A total of n = 5 mice per condition were analysed. Statistical significance was assessed by two-way ANOVA with Dunnett’s multiple comparisons test. Reported p-values correspond to endpoint tumour sizes on day 18. **(h)** Synergy index (SI) for inhibition of xenograft tumour growth by combined Trametinib and Silmitasertib treatment, calculated from the quantifications shown in (g). Error bars were calculated by error propagation using S.E.M. values. SI < 1 indicates synergy.

Based on this model, induction of rRNA synthesis downstream of oncogenic RAS signalling would be expected to be highly robust. Therefore, we hypothesised that effective suppression of ribosome biogenesis in PDAC may require simultaneous inhibition of both the initiating MAPK signal, as well as the CK2-dependent amplification loop, particularly in therapeutic settings where pathway inhibitions are often incomplete. To test this, we first assessed the effect of combined MAPK and CK2 inhibition on rRNA synthesis across decreasing inhibitor doses. Short-term Trametinib treatment at 5 and 10 nM concentrations potently blocked MAPK signalling, as shown by a complete loss of pErk1/2 signal, whereas 1 nM Trametinib produced only a partial inhibition (Supplementary Data Fig. S6a). Similarly, short-term Silmitasertib treatment at 5 and 10 µM strongly reduced CK2-dependent phosphorylations, whereas 1 µM Silmitasertib produced only a partial inhibition of downstream CK2 signalling (Supplementary Data Fig. S6b). Consistent with this, high-dose Trametinib or Silmitasertib treatments significantly inhibited rRNA synthesis, whereas the lowest doses of either inhibitor alone had no significant effect (Fig. 6b,c). Nevertheless, combining the lowest doses of Trametinib and Silmitasertib was sufficient to significantly suppress rRNA synthesis.

To assess whether this reflected synergy between the two inhibitors, we calculated a synergy index (SI) for rRNA synthesis inhibition based on Bliss independence analysis^39^. SI reveals existence of synergy between different treatments, with SI values < 1 indicating synergy, SI = 1 indicating additivity, and SI > 1 indicating sub-additivity. This analysis showed that Trametinib and Silmitasertib acted synergistically at their lowest combined doses, but not at higher doses, to inhibit rRNA synthesis (Fig. 6d). These findings support our proposed positive feedback model, in which complete inhibition of either MAPK or CK2 is sufficient to block rRNA synthesis. However, when each pathway is only partially inhibited, neither inhibitor alone is sufficient, and only their combined activity can disrupt the full signalling circuit to suppress rRNA synthesis.

We next assessed the effects of Trametinib and Silmitasertib on anchorage-independent growth of iKras cells using a 3D spheroid assay. This approach provides a more physiologically relevant *in vitro* model than 2D culture, including incomplete drug penetration that better resembles solid tumour tissue settings^40^. At the lowest doses tested, Trametinib or Silmitasertib alone did not significantly inhibit iKras spheroid growth, whereas combined treatment significantly reduced growth (Fig. 6e; Supplementary Data Fig. S6c). Higher-dose combinations were also more effective than either inhibitor alone for inhibiting spheroid growth, suggesting synergistic activity across the dose range tested (Fig. 6e; Supplementary Data Fig. S6c). Accordingly, SI analysis revealed strong synergistic inhibition of spheroid growth after longer treatment durations, at 9 and 12 days, across all tested drug doses (Fig. 6f). This delayed synergy is consistent with a mechanism that primarily targets ribosome biogenesis, as mature ribosomes are highly stable molecular assemblies with half-lives of several days^41^. Thus, reduced ribosome production would be expected to impair cell growth in a delayed manner.

Finally, we asked whether our *in vitro* spheroid-based findings extend to tumour growth *in vivo*. CD1 nude mice maintained on doxycycline-containing water were subcutaneously injected with iKras cells and allowed to form xenograft tumours for one week. Mice were then treated orally with Trametinib and Silmitasertib, either alone or in combination, and tumour growth was continuously monitored for 18 days. As controls, an untreated doxycycline-fed group, and an untreated doxycycline-withdrawn group were included. In agreement with previous studies^2,17^, Doxycycline withdrawal led to regression and eventual loss of tumours, further confirming the strict dependence of the iKras model on Kras^G12D^ expression. Importantly, whereas Trametinib or Silmitasertib alone failed to significantly inhibit endpoint tumour growth *in vivo*, their combined treatment effectively blocked tumour growth (Fig. 6g). SI analysis further revealed a strong synergistic inhibition of tumour growth by the combination treatment (Fig. 6h). Importantly, as in the 3D spheroid assays, the synergy emerged only at later treatment timepoints, consistent with a primary mechanism involving ribosome biogenesis suppression.

PDAC is characterised by a prominent desmoplastic stroma that can limit drug delivery to cancer cells^42^. We therefore also assessed endpoint tumour tissues by IHC to validate target engagements by Trametinib and Silmitasertib. CK2 showed strong nucleolar enrichment in untreated tumours, which was significantly reduced by Silmitasertib treatment, consistent with our finding that CK2 nucleolar localisation depends on its own kinase activity (Supplementary Data Fig. S6d,e). Trametinib also reduced nucleolar CK2 localisation, although to a lesser extent than Silmitasertib, consistent with the role of MAPK signalling in promoting rRNA synthesis upstream of CK2 (Supplementary Data Fig. S6d,e). We also assessed pErk1/2 levels within the tumours by IHC. Control tumours showed a strong pErk1/2 staining, which was significantly reduced by Trametinib, but not Silmitasertib treatment (Supplementary Data Fig. S6d,f). Together, these results demonstrate that orally administered Trametinib and Silmitasertib successfully reach PDAC tumour cells *in vivo* to inhibit MAPK and CK2 signalling, with their combined treatment leading to synergistic inhibition of PDAC tumour growth.

## Discussion

Prominent nucleoli were amongst the earliest recognised cytological features of malignant cells^5^, but mechanisms that link the nucleolus to oncogenic drivers of malignancy remain poorly understood. Here we show that oncogenic RAS signalling remodels the nucleolar form and function through a self-amplifying rRNA circuit. In PDAC cells, oncogenic RAS–MAPK activation promotes coalescence of smaller nucleoli into larger ones. This morphological transition is coupled to selective remodelling of nucleolar protein composition, phosphorylation landscape, and sub-compartment organisation, revealing that cancer-associated changes to nucleolar morphology are intimately linked with nucleolar function. Our quantitative spatial proteomic and phosphoproteomic analyses identify CK2 as a major kinase that is recruited to the nucleolus upon RAS–MAPK activation, where it phosphorylates many components of rDNA chromatin and the RNAPI transcription machinery. This leads to enhancement of rRNA synthesis, which is both necessary and sufficient for promoting nucleolar fusion. These findings suggest that the enlarged nucleoli that are characteristic of malignant cells arise, at least in part, from sustained nascent rRNA-driven coalescence of nucleolar material.

A key insight from this work is the notion that CK2 recruitment to the nucleolus is mediated by nascent pre-rRNA itself, adding to the growing body of evidence that suggest pre-rRNA is not simply a passive substrate during ribosome production, but actively organises this process in time and space^38,43,44^. Specifically, we reveal that the 3′ETS regions of pre-rRNA acts as a recruiting scaffold for CK2. As the end segment of 47S pre-rRNA, 3′ETS is only generated upon successful completion of RNAPI-mediated transcription, and is thought to be rapidly removed during the first stages of rRNA processing^36^. In yeast, processing of 3’ETS and the upstream ITS1 have been shown to be coupled^45^. However, little is currently known about the involvement of 3’ETS in coordinating specific steps of ribosome biogenesis in higher eukaryotes. We here reveal that through CK2 recruitment, this transient pre-rRNA spacer sequence functions as an active regulatory scaffold to reinforce rRNA synthesis through a positive feedback loop. Importantly, 3’ETS is ideally positioned for such a role, as its presence reports successful completion of rDNA transcription, thus licensing further transcriptional activation only if productive rRNA synthesis can be completed.

Our model also provides a mechanistic explanation for how pathological nucleolar enlargement specifically arises in cancer. In non-transformed cells, transient RAS–MAPK signalling would be expected to produce a temporary increase in rRNA transcription, with the short-lived 3’ETS-mediated feedback mechanism acting to extend rRNA synthesis in a transient manner to support physiological ribosome production. In malignant cells, however, constitutive RAS–MAPK signalling drives persistent 3′ETS production, leading to sustained CK2 recruitment to the nucleolus, and a runaway amplification of RNAPI transcription. This results in substantial accumulation of nascent pre-rRNA in the nucleolus, which in turn drives nucleolar fusion. Crucially, although defined here in the context of RAS–MAPK and PDAC, the 3′ETS–CK2-mediated positive feedback mechanism may similarly operate downstream of other oncogenic drivers of rRNA synthesis, such as MYC and mTOR, which are also known to directly activate rDNA transcription^46^, making this mechanism broadly applicable to diverse cancers. In support of this, we demonstrate that nucleolar enrichment of CK2 is a common feature of several solid tumours.

Finally, the positive feedback mechanism uncovered here has potential therapeutic implications. MAPK pathway inhibitors have shown limited efficacy in PDAC, despite the near-universal dependence of this disease on oncogenic RAS–MAPK signalling^42^. Our data suggest that these limitations may be overcome through the combined use of MAPK inhibitors with CK2 inhibitors, which together disrupt an otherwise robust ribosome biogenesis programme that is likely central to the promotion of tumour growth. We show that while partial inhibition of either MAPK or CK2 alone is insufficient to suppress rRNA synthesis, their combined inhibition can effectively block rRNA synthesis by interfering with both the MAPK-dependent initiating signal and the CK2-dependent amplification loop. This dual targeting synergistically suppresses rRNA synthesis, anchorage-independent 3D spheroid growth, and *in vivo* tumour growth in mouse xenografts. Whether this combination strategy will be effective in treating PDAC in patients remains to be determined.

In summary, we identify the 3′ETS region of pre-rRNA as a scaffold for recruiting CK2 to the nucleolus downstream of oncogenic RAS signalling. We show that this pre-rRNA-driven kinase localisation mechanism acts as a positive feedback loop to amplify rRNA synthesis, promote nucleolar fusion, and mediate PDAC cell proliferation and tumour growth. Our work reveals that nascent pre-rRNA actively shapes the morphology and function of the nucleolus through specific RNA–protein interactions, and provides a molecular mechanism for malignancy-associated hyperactivation of ribosome biogenesis and nucleolar enlargement, which was first reported 130 years ago^5^.

## Methods

List of antibodies and oligonucleotides used in this study can be found in Supplementary Tables S1 and S2.

### Cell culture

iKras PDAC cells^2^ were cultured in DMEM (Gibco) supplemented with 10% FBS, 1% penicillin/streptomycin, and doxycycline (1 μg/ml), in a humidified incubator at 37°C with 5% CO2. To remove Kras^G12D^ expression, cells were starved of doxycycline for 48 h. Re-expression was achieved by doxycycline addition for 24 h. For colony formation assays, cells were seeded at 1,000 cells per well, in triplicate for each experimental condition, with media and treatments replenished every 2-3 days. After 6 days, cells were washed with 1X PBS, stained with Crystal Violet (0.5% w/v Crystal Violet, 1% Formaldehyde, 1% MeOH in PBS) for 1 h, before being washed with water until desirable contrast was reached. Plates were then dried and imaged on an Amersham Imager 600. For 3D-spheroid assays, cells were seeded with doxycycline at 200-8000 cells per well in low attachment round-bottom tissue culture plates, and incubated for 3 days to allow for spheroid establishment. Spheroids were then subject to the indicated treatments for up to 12 days, with refreshment of media every 2 days. Brightfield images of each spheroid were taken on a Leica Thunder microscope, before cell viability was determined by CellTiter-Glo 3D (Promega), following manufacturer’s instructions.

### Plasmids, DNA transfections, and stable cell line generations

Wild-type and NIKx3-tagged CK2α constructs were synthesised by GeneArt (Thermo Fisher Scientific) into a Gateway pDONR221 entry clone, and were subsequently subcloned into the pcDNA™6.2/C-EmGFP-DEST destination vector, using the Gateway LR reaction kit (Thermo Fisher Scientific), according to the manufacturer’s instructions cloning (Invitrogen). Constructs were transfected into iKras cells, and the transfected cells were subjected to antibiotic selection with 5 μg/mL of blasticidin for 10 days. GFP-positive cells were then isolated by Fluorescence-Activated Cell Sorting (FACS) using a BD FACS Aria II. TAPIR PiggyBac mammalian expression construct containing rDNA promoter-targeting guide RNAs, dCas9 fused to VP64, RelA, and Rta transactivation domains, and a P2A-fused EGFP reporter^31^, was transfected into iKras cells, along with Super PiggyBac Transposase (Cambridge Bioscience), using Lipofectamine 2000 (Thermo Fisher Scientific) as per manufacturer’s protocol, followed by FACS sorting to enrich for GFP-positive cells.

### ASO and siRNA transfections

Cells were seeded at 50,000 cells per well of a 6-well tissue culture plate. siRNA-mediated knockdowns were performed using Dharmacon ON-TARGET plus siRNAs, as shown in the table S2, with Lipofectamine RNAiMAX, as per manufacturer’s instructions (Thermo Fisher Scientific). Cells were analysed after 72 h of incubation with the transfection mix and knockdown efficiency was determined by RT-qPCR and western blotting. For the transfection of ASOs, iKras cells starved of Doxycycline for 48 h were seeded onto iBidi 18 well μ-slides at 5,000 cells per well, with or without the addition of Doxycycline, and allowed to attach for 6 h. Tiling Antisense DNA oligonucleotides mapping the 5’ETS or 3’ETS regions of the *mus musculus* 47S pre-rRNA were transfected into the cells using Lipofectamine RNAiMAX, as per manufacturer’s instructions, and incubated overnight. At 18 h post-transfection, cells were subjected to nascent RNA imaging and immunofluorescence analysis as described below. Details of all ASO sequences can be found in Table S2.

### Immunofluorescence

Immunofluorescence was performed in iBidi 18 well μ-slides. 1,000 cells per well were seeded without doxycycline and incubated for 48 h, before addition of doxycycline or other indicated treatments. For the imaging of nascent RNA, 5-Fluorouridine (2 mM) was added to each well and incubated for 30 minutes to allow for pulse labelling of newly synthesised RNA. Cells were washed with PBS and fixed with 4% Formaldehyde in PBS for 15 minutes at room temperature (RT). Fixed cells were then permeabilised with 0.5% Triton-X100 in PBS for 10 minutes at RT, followed by three PBS washes. Cells were blocked in 4% BSA for 30 minutes, followed by incubation with the indicated primary antibodies (diluted in blocking buffer), either for 1 h at RT or overnight at 4°C. After primary antibody incubations, slides were washed with PBS three times, before incubation with fluorophore-conjugated secondary antibodies and Hoechst (diluted in blocking buffer) for 1 h at RT in the dark. Slides were then washed three times with PBS and imaged on a Zeiss LSM 770, Zeiss LSM 880 or Leica Thunder microscopes, using a 63X oil or water immersion lens. All buffers for nascent RNA imaging were made with RNase free reagents and the blocking buffer was supplemented with SUPERase-In RNase Inhibitor. For RNase treatment, 1 unit of RNase A/T1 mix (Thermo Fisher Scientific) per well was added to the blocking buffer and slides were incubated at RT for 1 h. In control conditions, SUPERase.In RNase Inhibitor was added instead at a 1:500 dilution. Image analysis was performed using ImageJ or FIJI, where individual nucleoli were identified by making a binary mask of the nucleolar marker (Ncl or Npm1) which was then converted to individual ROIs using the “analyse particles” function. The area and integrated intensity of the channel of interest was measured for each individual ROI and statistical analysis was performed in GraphPad Prism. Details of all antibodies used in immunofluorescence can be found in Table S1.

### Live cell imaging

10,000 iKras cells, starved of doxycycline for 48 h prior to the experiment, were seeded on 2 mL glass bottom individual microscopy chambers (Thermo Fisher Scientific) and allowed to attach overnight. The next day, cell chambers were mounted on an Evident ScanR widefield microscope equipped with a controlled incubation chamber maintained at 37°C in 5% CO_2_, and allowed to acclimatise to the chamber for 4 h, followed by 20 h of imaging, with or without doxycycline introduction. Images were collected every 10 min, using a phase contrast 100x, 1.45 NA lens at 100ms speed.

### Immunohistochemistry

IHC analysis of human tissue samples were performed on a cohort of 20 human pancreas tissues, with matched sections from PDAC, PanIN, and adjacent normal tissue regions, obtained from the Pancreatic Cancer Research Fund Tissue Bank (PCRFTB) as formalin fixed paraffin embedded tissue slides. Tumours from the iKras mouse xenograft experiments were prepared by Barts Cancer Institute Pathology Core Facility as formalin fixed paraffin embedded tissue slides. Tissue slides were deparaffinised with two 5-minute washes in xylene and rehydrated in 100% then 90% and 70% Ethanol for 2 minutes each. Endogenous peroxidase activity was blocked with a 10 minute incubation in 2% H_2_O_2_ in Methanol followed by washing in dH2O. Antigen retrieval was performed by submerging the slides in antigen retrieval buffer (10 mM Sodium Citrate in dH2O, adjusted to pH 6) and heating either in a pressure cooker for 10 minutes at 95°C or a microwave set to 50% power for 10 minutes. After allowing to cool, a hydrophobic barrier was drawn around the tissue. Slides were incubated in blocking buffer (5% Goat serum in 0.2% PBT-T) for 1h followed by overnight incubation in primary antibody diluted in SignalStain Antibody Diluent (Cell Signalling Technologies) at 4°C in a wet chamber. The next day, SignalStain Boost Detection HRP Reagent (Cell Signalling Technologies) was equilibrated to RT while the slides were washed 3 times with wash buffer (0.2% PBS-T). Two drops of the SignalStain Boost Detection HRP Reagent were added to each slide and incubated in a wet chamber for 1 h followed by three washes with wash buffer. One drop (∼30 μL) of SignalStain DAB Chromogen Concentrate (Cell Signalling Technologies) was diluted in 1 mL SignalStain DAB Diluent. To each slide, 100 μL of the DAB solution was added and incubated for 5-10 minutes at RT. Each slide was washed in dH2O for 5 minutes before clearing and mounting. Stained slides were dehydrated in 90% and 100% Ethanol and cleared in Xylene before mounting a cover glass slip with permanent mounting media (VectaMount). Images were acquired on either a NanoZoomer S210 slide scanner at 40X magnification, or an Olympus ScanR wide-field at 40X magnification. Details of all antibodies used for IHC can be found in Table S1.

### Western blotting

Samples were collected from tissue culture plates in 4% SDS, 100 mM Tris-HCl pH 7.5 sonicated with a probe sonicator (SoniPrep 150, MSE) at 50% power for 5 cycles of 10 seconds on and 10 seconds off. Sample protein concentration was estimated with a Pierce BCA Protein Assay Kit (Thermo Fisher) and adjusted with the same lysis buffer to balanced protein concentrations. NuPAGE 4x LDS Sample Buffer (Thermo Fisher) and reducing agent; DTT (100 mM), was added to each sample before denaturing by boiling at 95°C for 5 minutes. Samples were loaded onto a NuPAGE 4%–12% Bis/Tris protein gel (Thermo Fisher) and ran in NuPAGE MOPS buffer at a constant of 150 V for 60-90 minutes. Protein was transferred to an Immobilon-P PVDF membrane (Millipore) using a standard wet transfer device ran at a constant of 1 Amp for 2 h in transfer buffer (1X Tris Glycine, 20% methanol in dH2O). Membranes were blocked in blocking buffer (5% BSA in PBT-T) for 1 h, followed by primary antibody incubation (diluted in blocking buffer) overnight at 4°C. Membranes were washed three times in PBS-T, followed by incubation with anti-mouse or rabbit HRP-conjugated secondary antibodies, diluted in blocking buffer, for 1 h at RT. Membranes were then probed with Pierce ECL Plus HRP-detection reagent and developed and imaged on an Amersham Imager 600 (GE Healthcare). Details of all antibodies used for western blotting can be found in Table S1.

### Reverse Transcriptase quantitative Polymerase Chain Reaction (RT-qPCR)

Total RNA was extracted by TRIzol reagent (Thermo Fisher) as per manufacturer’s protocol. RT-qPCR was performed using Brilliant II SYBR Green One Step (Agilent) on an QuantStudio 5 instrument (Applied Biosystems). RNA expression levels were estimated using the ΔΔCT method, where the average change in cycle threshold of the target gene was normalised to the average change in cycle threshold of housekeeping genes. Details of all primers used for RT-qPCR can be found in Table S2.

### Nucleolar isolation

Nucleolar isolation was carried out as described previously^19^, with some modifications. Briefly, 1,000,000 iKras cells were seeded onto 10 cm dishes and grown for 48 h without doxycycline to remove Kras^G12D^. Cells were then treated with or without doxycycline (1 μg/ml), along with the indicated inhibitors for the stated timings. Plates were subsequently washed three times with ice-cold buffer S1 (0.5 M Sucrose, 3 mM MgCl2), before collection in 3 mL of the same buffer supplemented with protease and phosphatase inhibitor cocktail tablets (Thermo Fisher). The cell suspensions were sonicated on a probe sonicator at 50% power (10 mAmp) for 5 cycles of 10 seconds on and 10 seconds off. Each sample was checked under a light microscope to ensure successful rupture of plasma membrane. Sonicated cell suspensions were under laid with Buffer S2 (1 M Sucrose, 3 mM MgCl2, supplemented with protease and phosphatase inhibitor cocktails) before centrifuging at 1,800 g for 10 minutes at 4°C. The pellet, containing the isolated nucleoli, was solubilised in 4% SDS, 100 mM Tris-HCl pH 7.5 for downstream western blotting or mass spectrometry analysis.

### Proteomics sample preparation

Fractionated nucleoli and their corresponding whole cell lysates were prepared for LC-MS/MS analysis by isobaric Filter Aided Sample Preparation (iFASP) as described before^47^, with some modifications. Briefly, 60 µg of total protein from each sample was reduced with 10 mM Dithiothreitol (DTT) for 30 minutes, and alkylated with 55 mM iodoacetamide for 30 minutes in the dark. Samples were then diluted 7-fold in 8 M urea, 100 mM Tris/HCl pH 8.5 (UA buffer), and transferred to Vivacon 500 filters, before being concentrated by centrifugation at 14,000 g for 20 min. Filters were washed twice by serial addition of UA buffer followed by centrifugation. This was followed by three additional washes with 100 mM TEAB. Samples were subsequently digested by the addition Trypsin (1:100 w/w) in 100 µL of TEAB buffer to the filter tops and overnight incubation at 37°C. The following day, 300 µg of each TMTpro 18plex labelling reagent (Thermo Fisher) was added to each filter and incubated at 25°C for 1 h in a shaking thermomixer. Samples were then quenched by adding 5% hydroxylamine for 30 minutes. Peptides were eluted from filters into new collection tubes by centrifugation at 14,000 g. The elution was repeated twice with 100 µL of TEAB, and a final elution was carried out with 100 µL of 30% acetonitrile. Eluates with different TMT labels were then mixed together and dried with a vacuum concentrator. The pooled peptide mix split in a 1:9 ratio, where one part was fractionated into 7 fractions via Pierce™ High pH reverse-phase fractionation kit (Thermo Fisher), according to manufacturer’s instructions, and 9 parts were enriched for phospho-peptides using GL Sciences TiO_2_ phospho-enrichment kit, as per manufacturer’s protocol. Phospho and total fractions were then dried by vacuum centrifugation, before resuspension in A* buffer (0.1% TFA, 0.5% Acetic Acid, 2% Acetonitrile) for LC-MS/MS analysis.

### Targeted RNase H-mediated Extraction of crosslinked RBPs (TREX)

Proteins interacting with the 3’ETS region of rRNA in iKras cells were identified by a modified protocol of TREX^35^. Briefly, 800,000 iKras cells per replicate were seeded into 150mm tissue culture-treated dishes and grown for 48 h without doxycycline. Doxycycline was subsequently added to half of the dishes, and cells were incubated for a further 24 h, before being washed with ice-cold PBS and irradiated with 200 mJ per cm^2^ of UV-C (254 nm) on ice. Cells were then lysed in TRIzol reagent and combined into their respective treatment groups (Kras^G12D^ ON or OFF). Chloroform was added to each sample (200 µL for every 1 mL TRIzol) before homogenisation and centrifuging at 12,000 g for 15 minutes at 4°C. The upper aqueous and lower organic phases were removed from each sample, and the interphase was resolubilised in 1 mL TRIzol per 20 million cells. The phase separation was repeated four more times, and the final isolated interface was then washed gently with ddH_2_O to remove any traces of TRIzol, before being resolubilised and subjected to DNA digestion with TURBO DNase, as described before^35^. Samples were then precipitated by addition of 50% isopropanol, 150 mM NaCl and incubation at-20°C overnight, followed by centrifugation at 18,000g at 0°C for 15 minutes. The resulting pellet was washed with 75% ice-cold ethanol, before resuspension in 590 µL of Annealing buffer (100 mM Tris-HCl pH 7.0, 50 mM NaCl, 55% Formamide, in ddH_2_O). The resulting interface solution was then split into 10 x 54 µL aliquots. To each sample, 6 µL of a 100 µM pool of 60 nt-long tiling non-overlapping antisense DNA oligonucleotides complementary to the 3’ETS region of the Mus musculus 47S ribosomal RNA transcript was added. Samples were incubated at 50°C for 3 minutes to denature the RNA. For annealing of the antisense oligonucleotide probes and RNase H-mediated release of 3’ETS-crosslinked proteins, 180 µl RNase H digestion Mix (1 U/million cells Thermostable RNase H in 1X RNase H Buffer) was added to the ‘+RNase H’ samples, whilst the control ‘No RNase H’ samples were diluted identically but only with the 1X RNase H buffer. Dilution of formamide allowed for the annealing of the oligonucleotides to take place at 50°C. All samples were subsequently incubated for 1 h with shaking at 50°C to mediate RNase H digestion. Samples were then centrifuged at 12,000 g for 3 minutes at 4°C to remove any insoluble parts and the supernatant was transferred to a new tube. 10% of each sample was taken for RNA extraction and RT-qPCR analysis, whilst the remaining 90% was subjected to a final phase extraction using TRIzol LS to recover the released RBPs in the organic phase. The released RBPs were then precipitated by adding ice-cold acetone to a final concentration of 80%, and overnight incubation at-20°. The next day, samples were centrifuged at 16,000 g for 20 min at 4°C. The resulting pellet was washed 2 times with 80% acetone, followed by precipitation and centrifugation. The final protein pellet was air-dried and resuspended in 50 μL of 8 M Urea in 50 mM Ammonium Bicarbonate (ABC). Samples were reduced by DTT (10 mM) for 30 minutes at RT, and alkylated with IAA (55 mM) for 30 mins at RT in the dark. The urea was then diluted to 2 M by adding ABC, before 1 µg of Trypsin was added to each sample followed by overnight incubation at RT. Samples were desalted the next day using the C18 Stage-Tip procedure^48^, and recovered in A* buffer. 30% of each sample was used for TMTpro labelling (Thermo Fisher), as per manufacturer’s instructions. TMT-labelled samples were then pooled. As a carrier channel, 10 µg of trypsin-digested and TMT-labelled total iKras nucleolar fraction was also included in the mix. The combined peptide mix was then fractionated using Pierce™ High pH reverse-phase fractionation kit (Thermo Fisher), according to manufacturer’s instructions, before being resuspended in A* buffer for LC-MS/MS analysis. Details of TREX antisense oligonucleotides can be found in Table S2.

### Mass spectrometry

All LC-MS/MS analyses were performed on a Q Exactive-plus Orbitrap mass spectrometer coupled with a nanoflow ultimate 3000 RSL nano HPLC platform (Thermo Fisher). For total proteomics, equivalent of 1 μg of peptides resuspended in A* buffer was injected into the nanoflow HPLC. For phospho-proteomics and TREX analysis, ∼90% of the available peptide mixture was injected. Samples were resolved at flow rate of 250 nL minute−1 on an Easy-Spray 50 cm × 75 μm RSLC C18 column (Thermo Fisher), using a 123-minute gradient of 3% to 35 % of Buffer B (0.1% formic acid in acetonitrile) against Buffer A (0.1% formic acid in LC–MS gradient water). LC-separated samples were infused into the MS by electrospray ionization. Spray voltage was set at 1.95 kV, and capillary temperature was set to 255°C. MS was operated in data dependent positive mode, where one MS scan was followed by 15 MS2 scans (top 15 method). Full scan survey spectra (m/z 375–1,500) were acquired with a 70,000 resolution for MS scans and 35,000 for the MS2 scans. A 30 second dynamic exclusion was applied to all runs. Raw data files were searched against a mouse UniProt database using MaxQuant (version 2.2.0.0 or 2.6.5.0)^49^. For nucleolar proteome and phospho-proteome profiling, parameter groups were set to define the phospho-or total-proteome analysis and fractions were set for the total proteomics analysis. “Reporter ion MS2” type was selected, choosing 18-Plex TMT as the label type. Batch-specific TMT correction factors, determined by the manufacturer, were inputted, with a reporter mass tolerance of 0.003 Da. Enzyme specificity was set to “Trypsin,” allowing up to two missed cleavages. Phosphorylation (STY) was added to variable modifications for the phospho-parameter group. False discovery rates (FDR) were calculated using a reverse database search approach and were set at 1%. “Re-quantify” option was enabled. Default MaxQuant settings were used for all other parameters. For TMT-labelled TREX analyses, fractions were set for each relevant sample. For group specific parameters, ‘Reporter ion MS2” type was selected, choosing 18-Plex TMT as the label type, removing the unused labels. Batch-specific TMT correction factors, determined by the manufacturer, were inputted, with a reporter mass tolerance of 0.003 Da. Default modifications were used. Digestion mode was set as ‘Trypsin’ with 2 maximum missed cleavages. For global parameters, ‘match between runs’ was enabled, and an FDR of 1% was applied using a reverse database approach. “Re-quantify” option was enabled. All other settings were set as default.

### *In vivo* tumour growth analysis

Subcutaneous xenograft tumour growth experiments were performed under UK Home Office Project Licence PP6460882. A total of 1,000,000 iKras PDAC cells suspended in 100 μL Opti-MEM (Thermo Fisher) were injected into the lower flank of 5-week-old female CD1 nude mice. To maintain Kras^G12D^ expression, mice were provided with doxycycline (2 g/L) and sucrose (20 g/L) in their drinking water starting 3 days prior to injection. Tumours were allowed to establish for 1 week in presence of doxycycline, before treatment with Silmitasertib (75 mg/kg), Trametinib (1 mg/kg), or the combination, administered daily by oral gavage. Positive and negative control groups were maintained on doxycycline alone (no drugs) or had doxycycline withdrawn from the water. Treatments were continued for 18 days, and tumour volumes were measured twice weekly with callipers. Five mice per treatment cohort were used.

Orthotopic xenograft tumour growth experiments were performed under UK Home Office Project Licence PP9448177. 500,000 iKras PDAC cells, resuspended in 20 μL of 50% Matrigel (BD Biosciences) in Hanks’ balanced salt solution were injected into the pancreas of 5-week-old male CD1 nude mice. Surgery was performed under isoflurane anaesthesia, and animals received subcutaneous Vetergesic (buprenorphine) analgesia in the scruff before surgery, later the same day, and again the following morning. To maintain Kras^G12D^ expression, mice were provided with doxycycline (2 g/L) and sucrose (20 g/L) in their drinking water starting 3 days prior to orthotopic engraftments. Tumours were established for 2 weeks in the presence of doxycycline, before doxycycline was removed from water for half of the mice, with animals maintained for a further period of 2 weeks. *In vivo* tumour imaging was performed at 2, 3, and 4 weeks timepoints, using a 3 min T2-weighted MRI scan on a Bruker ICON 1T MRI system under anaesthesia with isoflurane. Three mice per cohort were used.

## Statistical analysis

All proteomics data analyses and visualisations were performed in Perseus (versions 1.6.2.1 or 1.6.6.3)^50^. Principal Component Analysis (PCA) was used to reveal the overall similarity of replicates. Any clear sample outliers revealed by the PCA were removed from downstream analyses. For 2-sample t-test analysis and multiple-sample ANOVA analysis, a permutation-based FDR calculation with an S0 of 0.1 was used. Category enrichment analysis was performed using Fisher’s exact test, with Benjamini-Hochberg FDR calculation. Synergy Index (SI) between Trametinib and Silmitasertib was calculated based on Bliss independent analysis, using the formula:

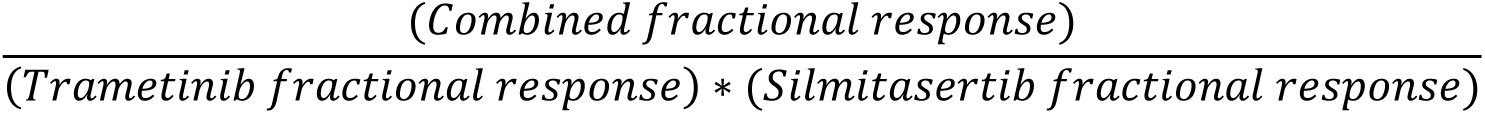

All other statistical analyses were performed in GraphPad Prism (version 11.0.1). Data were first tested for normality using Shapiro-Wilk normality testing. For significance testing between two conditions, either an unpaired t-test for normally distributed data, or a Mann-Whitney t-test for non-normally distributed data, was used. For significance testing between more than two conditions, either an ordinary one-way ANOVA for normally distributed samples, or a Kruskal-Wallis one-way ANOVA for non-normally distributed samples, were used. For significance testing involving two variables (e.g. treatment and timepoint) two-way ANOVA with repeated measures was used. Multiple comparisons correction was applied to all ANOVA analyses.

## Data availability

All mass spectrometry raw files and their associated MaxQuant output files were deposited on ProteomeXchange Consortium via the PRIDE partner repository, under the accession numbers PXD078625, PXD078637, and PXD078679.

## Author contributions

F.K.M. conceived the study, acquired the funding, and supervised the work. F.K.M. and E.L.A. wrote the manuscript. M.D. carried out the mouse orthotopic tumour growth analysis, and assisted with the 3’ETS TREX analysis. S.N.R. and M.D. conducted the xenograft tumour growth experiment, supervised by E.O and K.I.J. The nucleolar live-cell imaging analysis was performed by Z.R. The TAPIR system was generated by M.W. and S.H.S. The 3D confocal imaging of iKras colonies were performed by M.C. and Z.P., supervised by M.R.C and F.K.M. M.S.A. assisted with 3D Spheroid analysis. E.L.A performed all other experiments and data analyses.

## Acknowledgements

This study was funded by a Cancer Research UK (CRUK) Programme Foundation Award (DRCPFA-Nov24/100005), and a Biotechnology and Biological Sciences Research Council (BBSRC) project grant (BB/X007820/1) to F.K.M. E.L.A was supported by a Barts Cancer Institute (BCI) CRUK PhD studentship (C355/A25137). We acknowledge the BCI mass spectrometry, microscopy, and histopathology core facilities for supporting the MS, microscopy, and IHC experiments. We also acknowledge Oxford Micron Bioimaging Facility for supporting microscopy experiments. Human matched PDAC and normal tissue sections were provided by Pancreatic Cancer Research Fund Tissue Bank (PCRFTB), funded by the Pancreatic Cancer Research Fund. We would like to thank all members of the Mardakheh lab for their critical evaluation of the manuscript. We would also like to thank Vinothini Rajeeve, Tommy Shields, Sam Wallis, Hemant Kocher, Rhiannon Roberts, and Nadia Rahman for their assistance with sample collections and experiments.

