## Supplementary Information for "A pre-rRNA positive feedback loop drives malignant ribosome biogenesis"

Supplementary information for this manuscript consists of Supplementary Tables S1 and S2, Supplementary Data Figures S1 to S6, and a supplementary excel file containing Supplementary Datasets S1 to S15.

### Supplementary Tables

**Table S1: List of antibodies used in this study.**

| Antibody target | Host species | Company (catalogue number) | Dilution |
|---|---|---|---|
| **Immunofluorescence** | | | |
| **BrdU** | Mouse | Merck (B2531) | 1:500 |
| **CK2β** | Mouse | Santa Cruz (SC12739) | 1:50 |
| **CK2 phospho-substrates** | Rabbit | Cell Signalling Technology (8738S) | 1:400 |
| **Fibrillarin** | Chicken | Novus (NBP246881) | 1:100 |
| **Nucleolin** | Rabbit | Abcam (AB22758) | 1:400 |
| **Nucleophosmin** | Mouse | Invitrogen (32-5200) | 1:400 |
| **Alexa Fluor 488-conjugated Donkey** | Mouse | Jackson ImmunoResearch (715-545-150) | 1:1000 |
| **Alexa Fluor 594-conjugated Donkey** | Mouse | Jackson ImmunoResearch (715-585-150) | 1:1000 |
| **Alexa Fluor 647-conjugated Donkey** | Rabbit | Jackson ImmunoResearch (711-605-152) | 1:1000 |
| **Alexa Fluor 488-conjugated Donkey** | Chicken | Jackson ImmunoResearch (103-545-155) | 1:1000 |
| **Hoechst** | - | Thermo Fisher (H3569) | 1:2000 |

| Immunohistochemistry | | | |
|---|---|---|---|
| **CK2 β** | Mouse | Santa Cruz (SC12739) | 1:50 |
| **Phospho-MAPK (ERK1/2) p44/42** | Rabbit | Cell Signalling Technology (4370) | 1:500 |
| **Western Blotting** | | | |
| **CK2 α/ α'** | Rabbit | Cell Signalling Technology (2656) | 1:1000 |
| **GAPDH** | Mouse | Novus Biologicals (NB300-221) | 1:1000 |
| **KRAS (G12D mutant specific)** | Rabbit | Cell Signalling Technology (14429) | 1:1000 |
| **NPM1** | Mouse | Thermo Fisher Scientific (32-5200) | 1:1000 |
| **Nucleolin** | Rabbit | Abcam (22758) | 1:1000 |
| **Phospho MAPK (ERK 1/2) p44/42** | Rabbit | Cell Signalling Technology (4370) | 1:1000 |
| **Phospho-CK2 Substrate** | Rabbit | Cell Signalling Technology (8738) | 1:1000 |
| **Total MAPK (ERK 1/2)** | Rabbit | Cell Signalling Technology (9102) | 1:1000 |
| **α-Tubulin** | Mouse | Cell Signalling Technology (3873) | 1:1000 |

**Table S2: List of all oligonucleotides used in this study.**

| Oligo ID | Sequence (5'-3') | Company |
|---|---|---|
| **siRNAs** | | |
| **siCsnk2a1** | CCGAAGAGCCCUUUAAAUA, GGUCAGGGUUUACAGAGUA, CUGAACGAAUCAUGUCUUA, UCACCUGGCAUCAUAGAUA. | Horizon Discovery |

| | | |
|---|---|---|
| **siCsnk2a2** | GGUGGAACAAAUAUCAUUA,<br>UGACCAGCUUGUUCGAAUU,<br>GAACUAUAUGGUUAUCUGA,<br>GGAGUACAAUGUUCGAGUG. | Horizon Discovery |
| **siCsnk2b** | CCAAGUGCAUGGACGUGUA,<br>UCGCACAAAUGUUGGAAAA,<br>UGAGAUGCUUUAUGGGUUG,<br>UCUUAGACCUGGAACCUGA. | Horizon Discovery |
| **siURB1** | UUAAGUGGGUUCCGAGACA,<br>GCUCGAUGUUGUCCGCAAU,<br>AGGACAAGACUGCGAGAGU,<br>GUGCUGAGUUAUCGGGCCA. | Horizon Discovery |
| **siNT (ON-Target plus Non-Targeting Control)** | UGGUUUACAUGUCGACUAA,<br>UGGUUUACAUGUUGUGUGA,<br>UGGUUUACAUGUUUUCUGA,<br>UGGUUUACAUGUUUUCCUA. | Horizon Discovery |
| **Antisense Oligonucleotides** | | |
| **3'ETS_1** | TGGAAGAGGCGCCGCGCCGGACCCTCTCCGCCTCCACA<br>CCGCGACATCCTCCCCTCGTAC | Sigma |
| **3'ETS_2** | GCACGTCGCCGCCGCGGCGCGCGAAGAGGAGGGGCGG<br>GAGTGGGGATGCTCGGGCGGAGA | Sigma |
| **3'ETS_3** | GAGCGCCCCCCCGGCCGCCAGAGGCCTGGCGCCGAACG<br>CGGCGTTCACGGGAAAAACCCC | Sigma |
| **3'ETS_4** | TCCCGCCCCTCCAACCGGCCGTCCCCGACGGACGCGCC<br>GAGGATGGGGATCCCACCGTGC | Sigma |
| **3'ETS_5** | GTCACCGCGCCGCGGTCGCCGCCGCCGCCGATCCCCCT<br>CCTCCGCCCTCCGGCTTGGCCA | Sigma |
| **3'ETS_6** | AGGCCGGAGGGCAAGGGGCTGCGGCGGCAGGCGACGG<br>CAGGCCGCGGTCGCGACGGCACG | Sigma |
| **3'ETS_7** | AGGCTCCCACACCCAAGGAGAGCGACGGAAGGGGAAA<br>GAGAAACGAACCGTCCTCGGGAG | Sigma |
| **3'ETS_8** | ACGGAGGAGAAGACGATCCCCCCCCCAACCACCACACG<br>GTCGTTCGGGCAACCGCGCGGG | Sigma |
| **3'ETS_9** | GGCCGAGGCGACCGCCCCCGTGACGCCCGGGCGGCGAC<br>ACGGTCGCGTGCGAGCGAGCGGGCGTGCGACGG | Sigma |
| **5'ETS_1** | AAGGAAGCGCGAGCGGAGCGTCGGGGGTGGGACAGGA<br>CCAGGCGGGACAGAGAGCGCGAG | Sigma |
| 5'ETS_2 | AGAGGAGGAGGAGGCGGCGGCGGCGGAGAGCCTCGCG<br>GGAGGACAAACCGGGGGTGAGCC | Sigma |
| 5'ETS_3 | GGGCGGGCGCTCGCGCGGAACGGGAGAGTGCATGCGG<br>CCGGACCGACCCCGTCGGGGTCC | Sigma |
| 5'ETS_4 | GACAAAACCCGTCCGCGAGCACACGCACGCGGCAAAG<br>CCAACCCGAGCGTCGCCCCGGGA | Sigma |
| 5'ETS_5 | CAGCTCTCCGACACCTCTCTTACCCGCTCTCCCCTCCG<br>TCCCCGGCACAAATCCCGAACC | Sigma |
| 5'ETS_6 | CCTCGGCATCGGGAGAGCCGGAACGTCGACCGCACCG<br>GGCCGGGAGAGCAGGGTGGTGGG | Sigma |

| | | |
|---|---|---|
| 5’ETS_7 | TTGGCGTGGGGGTTGGCGGAAAGAGAAGCGCGACACC<br>GGCGAGACAGTCAAACCACGCGG | Sigma |
| 5’ETS_8 | GAGGGAATGACCGGTAGCACCATTCACTCCCCACACCA<br>ACGACCACCACCGGGACGGGCA | Sigma |
| 5’ETS_9 | GGCGGGGGTCGGCGGGGGCGAGCCCTGGATAGGCCAC<br>CCGTCGGAGCCCGGCGACACCCG | Sigma |
| 5’ETS_10 | CGTCTAGCCGCCGCGAGGCGGACACGCTCCCGACGCGA<br>CACAACGCCAGGGCCCAACTGA | Sigma |
| 5’ETS_11 | CCGCGGCGCGAAACCGCACACCGGCGGGTGCGGCAGG<br>GGCTCACGGGCGGGCGGAGGCCG | Sigma |
| 5’ETS_12 | CAGCGGGCCCCCCCCAAGCACACAGCAGCGAGGCGCA<br>ACGTCGGGGAACCGGGAAGCCCC | Sigma |
| 5’ETS_13 | CTGACCCTCCGGGCGAAGCCAGGAGTCCGCGAAGCCG<br>GCGAGAGAAGACCACGCCAACGG | Sigma |
| 5’ETS_14 | GGGTCGGGCAGGAGAGACCGGACAGCGGCCAGCCCCC<br>GGCGAGAAGCGGAGGCGGAGGAC | Sigma |
| 5’ETS_15 | GAACCCACCGTGAACTCGGGCGGCGCCATTGCGCGAG<br>AGATGGAAGGACGGAGGAGGGGA | Sigma |
| 5’ETS_16 | GGTCGACGAGCGACTTGAACCCCGCGAGAGGGAGGAA<br>CGAGGGGAGGGCCGAGAGCCCGC | Sigma |
| 5’ETS_17 | GCGGAGAAGGCCTGAGACCTCGGGGAGAGGGAGGTAG<br>ACATCGACCGGAGCCCCCACCGC | Sigma |
| 5’ETS_18 | CACACCGGGACACGCGCGCGGGGTTAATCCCACCGCCA<br>GGGTACCCACCTCGGCACTCCG | Sigma |
| 5’ETS_19 | GCAGGCCCCAACACGCGCCTTCCCGACGGACGACAGG<br>GCCGGACGAGCCCCCCAGACGCA | Sigma |
| 5’ETS_20 | CCTCGGCCAACAGACCACCACACCGTCGGGACAATGAC<br>CACTGCTAGCCTCTTTCCCTTT | Sigma |
| 5’ETS_21 | GCCCTCGCGGGGCAACCTTTTCCCCTCGCACGCAGCCC<br>TCTCCAAGACCGCGCAAGACAC | Sigma |
| 5’ETS_22 | CGCCACGGCCGCGGGGCCGGGACGCCCTCCAAACCAC<br>AGCTCCTCCCGCCCGCCGAGCCG | Sigma |
| 5’ETS_23 | ACTTTCTTACCCCTCGGCCCCCCCCCAAGACCCCGGTG<br>ACCGCCCGGACGCGTACGTGCC | Sigma |
| 5’ETS_24 | CGGCCGCCCCCATGCGGGGGGTCCGACAGCACGGCCAC<br>GGGTCAGTCAGAGGAGAGGGGG | Sigma |
| 5’ETS_25 | GGAGAGAGGAAAAAAGACCGGCTGCGGCGAGCGGGG<br>AGGGGGGAACGACACAGCAGAACG | Sigma |
| 5’ETS_26 | AGGGCCGGCGACGACGGCAAGGGCGGCCGAAACGGGC<br>CCCTCCCGGAAGGGGGTCCTCAC | Sigma |
| 5’ETS_27 | AAGGGACCGAGAGACCGCTCGGGCGTCGGACCGGTCA<br>GGGCACGCGAGGCAAAACCGGCG | Sigma |
| 5’ETS_28 | GGAATCACACTGCACCACTCCCCTGAGCAGTCCCACCA<br>CACCTGCAGCGCCGAGCCGACC | Sigma |
| 5’ETS_29 | CGAAACACGACGGCCCCCCGGGAGCCCGAGGAGGCGA<br>CACCGACACGAGAGAGAGACCGA | Sigma |
| 5’ETS_30 | TGCCGACACACCGATGCCTACCGACGAAGACCCCCGAG<br>CGGCAGTGGGTCCCCACGTCCG | Sigma |

| | | |
|---|---|---|
| 5'ETS_31 | ACGTCGCGCCGGACGCGGCCAACAGCCGTTCTCCCACA GAACGGTCCCGTCCCCGAGAGC | Sigma |
| 5'ETS_32 | GAGATCAATCAACGCCAGACACGGACCCTCTCGACCCC GAGGAGGGGAAGGAGGACTCCC | Sigma |
| 5'ETS_33 | GCGATTTCGCCAGCCGTGGGAACGCTGCCGCGGGAAG GGGGGGGCGAGGCGACACAACCA | Sigma |
| 5'ETS_34 | CACAGCCCGAGAAGCACACAGCACGCACCACGCGCGG CGCAGGCCGCCGTCGCGGGAGCG | Sigma |
| 5'ETS_35 | GCCAAAGTCGTGGACGCGAGCGAGAGGGACGCACGCG ACGGAAGGCACGGGTGAGGCACG | Sigma |
| 5'ETS_36 | GGAAGACGGGTGGGGACCCGGGAGCCCACCGTCGCCT ACCCACACCCCGTCCGCACGGCG | Sigma |
| 5'ETS_37 | GCCCTCAGGAGGGGAGGAAAGTACGCGCACGCGGGGG TGTGCACGGAGCCCTCTCCCCAG | Sigma |
| 5'ETS_38 | CCACCAACCACACAAGCCGAGCCACATGCTCCGCAGCA ACGGCAGGACGACAGACAGGCT | Sigma |
| 5'ETS_39 | CTGCCCCGCGTGATCCCTCCCCGAACTCGGAGCGGGGA GGCCGCGGGCCGCGGTAGACGA | Sigma |
| 5'ETS_40 | GAGAGCAAACGCTGCGCGCGCGGGAGGCGGTCAGGGG GCGAGCCCTCCAGGGGACCATTC | Sigma |
| 5'ETS_41 | GGGAGAGACCCCTCGAGACCGTAAGAAGCCCGCACCC TTTCCGGAAGATAGCTAGAGAAG | Sigma |
| 5'ETS_42 | GAAACTTTCTCACTGAGGGCGGGACGGACCCAAAACC CCGACGAGCCCCGGCCACCGAGA | Sigma |
| 5'ETS_43 | GAGGATGCATGCGACGAGCACACAGGGAAACCAGAAG ACCAACACGCGGGCGACCGCGCG | Sigma |
| 5'ETS_44 | GGGGGGTCTGCGGCAACAGGAAGGTTCTCTTCCAAGG GCATTCTGAGCATCCGCGGACAG | Sigma |
| 5'ETS_45 | GGGAGACCCTGCAACCGGTCCCCCCCGAAAGCCAGGC CTCTCAAAGCTCCCCACGGGAAA | Sigma |
| 5'ETS_46 | GCAATGAGTCTCTCGAGAAACTTTCCAACCCCAGCCG CGACCGCTCAGCACCAGCCCTAC | Sigma |
| 5'ETS_47 | GCACAGGCGTACTAGTCGCGGGGTGGGCGCCTCACCCG ACGAAGAAGGGGGGTCTTGCCC | Sigma |
| 5'ETS_48 | CTCCTTCTCTCCTCGACCCCCCTCTCACGGGCTTCTCA GACACAAACGGGAAGGCACACA | Sigma |
| 5'ETS_49 | GCCAGACGGAGCACCGGAGGCACACCAGGGAATGGGG AGCGCCACCACCAGCTCAGAAGC | Sigma |
| 5'ETS_50 | AGGCACCTAGGAGACAAACCTGGAACGCTCCAGGAGC ACCTCGACGCTTACAAGAAACAG | Sigma |
| 5'ETS_51 | CGCGTGCACACACCAGACGGGAAGGGTATGCAACGCC ACCGGCCACATCCACCGACTCTG | Sigma |
| 5'ETS_52 | GGAACATGGTCAAGCGAGACACGACCAAAGTGAAACA CGTGAGGGCACAACCGGGCGCCT | Sigma |
| 5'ETS_53 | CACACATCCACAAGGACCACGCGAACCACTGAGAAAA GTGCGCGCGGGGCAGCGTCGGGC | Sigma |
| 5'ETS_54 | CGGAGAGCCGCACCCCTGCCTTCCACACACCACCGGCA GACGGATGGGGTGGAGAGACGA | Sigma |

| | | |
|---|---|---|
| 5'ETS_55 | GGGCCCGCTGGCAGAACGAGAAGAGCGGCCGCCATTC GCCATGAATGTCCGTCCCTCGCC | Sigma |
| 5'ETS_56 | TGGCGCGGCTTGGCCCTGGCCCGAAGAGAACTCCGGA GCACCACATCGATCTAAGAGTGA | Sigma |
| 5'ETS_57 | GCAACGACGCGCAATCGGGAGAAACAAGCGAGATAGG AATGTCTTACACGCGGGGCAAGA | Sigma |
| 5'ETS_58 | CAGTTACTGATACGGGCAGACACAGAACAAGAGAACA CAACGAGCGACTGCCACAAAAAA | Sigma |
| 5'ETS_59 | AAGTGCACTCGGGAGGCACGTGGCATGAACACTTGGA CACCACAGACAGGAGTGAAGTAC | Sigma |
| 5'ETS_60 | TCGGGACTCTCCACCTCCCCAAAAAAAAAAAAAAAAA AAGAAAAAAAACAAAAAGAAATG | Sigma |
| 5'ETS_61 | CACTCGGGAGGCATGTGGAAGAAAGACCGGGAAGAGA AAAGAGCGGAGGTTCGGGACTCC | Sigma |
| 5'ETS_62 | AAGAGAAAAAAAAAAAAAAAAAGTGAGCCGAAATAA GGTGGCCCTCAACCACAGCTGGCT | Sigma |
| 5'ETS_63 | CCACCATTCCAACCGGGACAGGTGTGACAACGACACC TCGGGGAAATCGGGAAAAACGTC | Sigma |
| 5'ETS_64 | TGACACGCAGCAAAGTCACAGCGCCCAGTTGTCACAA GCTGCCCACAGCAAGCACACGCA | Sigma |
| 5'ETS_65 | AGCAGCAAGCCCGGGAAAGCAGGAAGCGTGGCTCGGG GACAGCTTCAGGCACCGCGACAG | Sigma |
| 5'ETS_66 | ACCCAAGCCAGTAAAAAGAATAGGCTGGACAAGCAAA ACAGCCTTAAATCGAAAGGGTCT | Sigma |
| 5'ETS_67 | CTTTATAGTGTCCTTTAGTGTTAATAGGGAAAGGACAG CGTGTCAGT | Sigma |
| **qPCR primers** | | |
| **Csnk2a1** | Mm_Csnk2a1_2_SG | QIAGEN (QT01767724) |
| **Csnk2a2** | Mm_Csnk2a2_va.1_SG | QIAGEN (QT01564066) |
| **Csnk2b** | Mm_Csnk2b_1_SG | QIAGEN (QT00173397) |
| **Urb1** | Mm_Urb1_2_SG | QIAGEN (QT01162693) |
| **Gapdh** | Mm_Gapdh_3_SG | QIAGEN (QT01658692) |
| **3'ETS rRNA FWD** | GGATCGTCTTCTCCTCCGTC | Sigma |
| **3'ETS rRNA RVS** | CGACGGAAGGGGAAAGAGAAA | Sigma |
| **Actin FWD** | GATGTATGAAGGCTTTGGTC | Sigma |
| **Actin RVS** | TGTGCACTTTTATTGGTCTC | Sigma |
| **18S rRNA FWD** | CAGTTATGGTTCCTTTGGTC | Sigma |
| **18S rRNA RVS** | TTATCTAGAGTCACCAAGCC | Sigma |

### Supplementary Figures

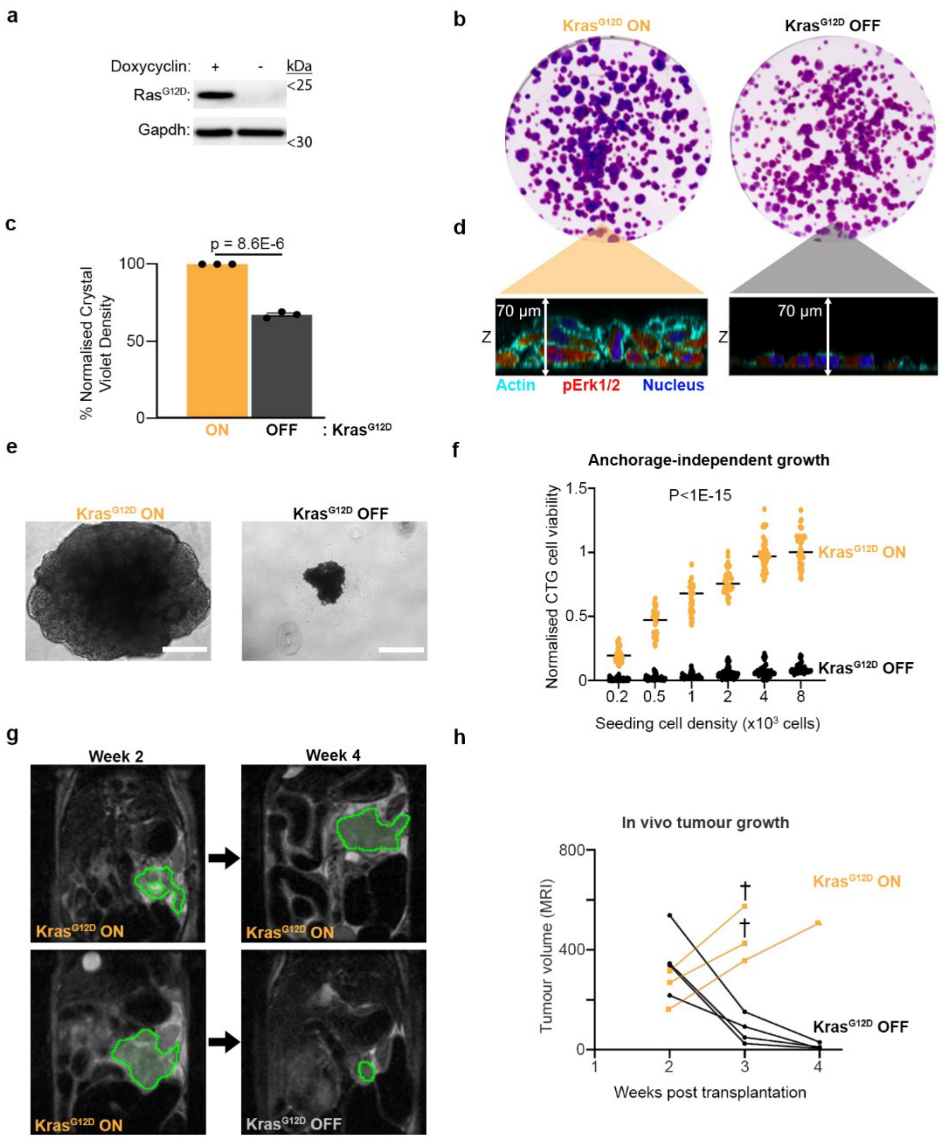


**Supplementary Data Figure S1: iKras provides an inducible isogenic model of malignancy.**

**(a)** Western blot analysis of $Kras^{G12D}$ expression in iKras cells treated with or without doxycycline for 24 h. $Kras^{G12D}$ was detected using a mutation-specific $RAS^{G12D}$ antibody. Gapdh was used as a loading control.

**(b)** Colony formation assay of iKras cells cultured for 6 days with or without doxycycline to induce $Kras^{G12D}$ expression. Colonies were visualised by Crystal Violet staining. Although iKras cells proliferate in the absence of $Kras^{G12D}$, induction of $Kras^{G12D}$ results in dense, multilayered cell colonies.

**(c)** Quantification of Crystal Violet staining intensity from the assay shown in (b). A total of n = 3 independent biological replicates were analysed. Significance was assessed using a two-tailed unpaired t-test.

**(d)** Z-projection confocal images of iKras cells cultured with or without doxycycline for 5 days. $Kras^{G12D}$ expression promotes the formation of multilayered cellular foci. Cells were immunostained for phosphorylated Erk1/2 (red), the key downstream effector of $Kras^{G12D}$ signalling. F-actin was visualised with phalloidin (cyan), and DNA was stained with Hoechst (blue).

**(e)** Representative phase-contrast images of iKras 3D spheroids cultured in low-attachment plates with or without doxycycline. 8,000 cells were seeded and cultured for 6 days before imaging. $Kras^{G12D}$ expression promotes anchorage-independent growth. Scale bar, 200 μm.
**(f)** Quantification of spheroid viability from (e). Cells were seeded at indicated densities with or without doxycycline and viability was measured using the CellTiter-Glo 3D assay after 6 days. Values were subsequently normalised to the maximal measured CTG value. A total of n = 40 independently seeded spheroids were analysed per condition. Significance was assessed using a two-tailed unpaired t-test.
**(g)** Representative MRI images of orthotopically established iKras tumours in nude mice. A total of $5 \times 10^5$ iKras cells were injected into the pancreas of CD1 nude mice, and tumours were established by administering doxycycline for 2 weeks. Animals were then divided into two groups: one remained on doxycycline, whereas doxycycline was withdrawn from the second group to switch off $Kras^{G12D}$ expression. Mice were imaged by MRI at weekly intervals. Representative tumours (green) from the same animals at weeks 2 and 4 are shown.
**(h)** Quantification of tumour volumes from the MRI analyses shown in (g). Each line represents an individual animal. Continued doxycycline administration resulted in progressive tumour growth, whereas doxycycline withdrawal led to tumour regression. † indicates animals that were euthanised before the endpoint owing to disease burden.

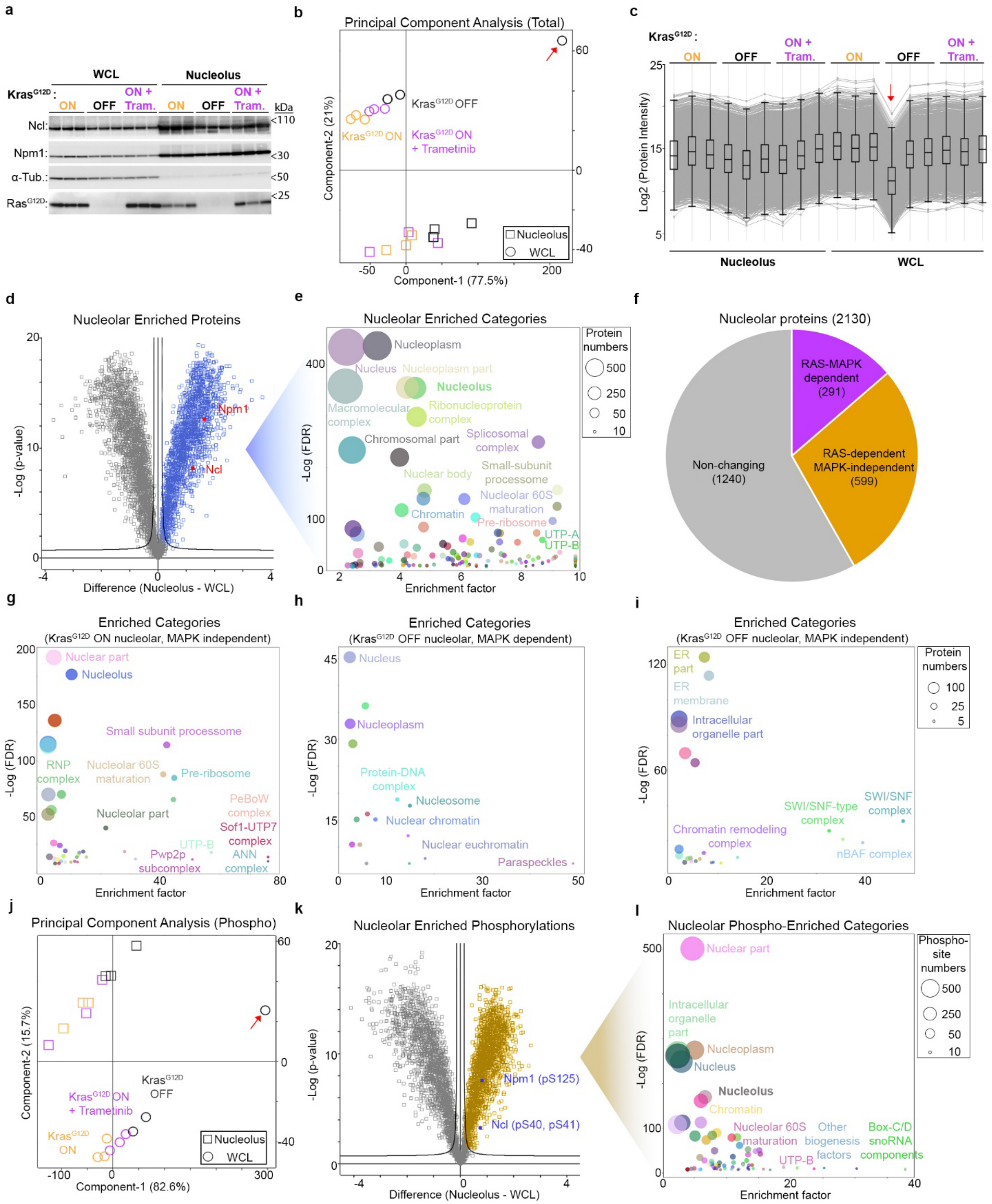


**Supplementary Data Figure S2: Characterisation of nucleolar proteome and phosphoproteome in iKras PDAC cells.**

**(a)** Western blot analysis of nucleolar fractions and matched whole-cell lysates (WCLs) from iKras cells treated with or without doxycycline for 24 h to induce $Kras^{G12D}$ expression, or with doxycycline plus Trametinib (10 nM) co-treatment to inhibit RAS–MAPK signalling. Nucleolin (Ncl) and nucleophosmin

(Npm1) were used as nucleolar markers, while α-Tubulin was used as a cytoplasmic marker. $Kras^{G12D}$ was detected using a mutation-specific $RAS^{G12D}$ antibody. Three independent biological replicates were analysed per condition.

**(b)** Principal component analysis (PCA) total proteomic profiles obtained from nucleolar fractions and their matched WCLs of iKras cells, under the indicated conditions. Samples show a clear separation between nucleolar and WCL, as well as an overall clustering of biological replicates within different treatment group. The red arrow indicates an outlier WCL sample which was removed from subsequent analysis.

**(c)** Protein intensity profile plots corresponding to the proteomic dataset shown in (b). The red arrow indicates the outlier WCL sample also highlighted in (b).

**(d)** Volcano plot comparing all nucleolar fractions with matched WCL samples from (b). Statistical significance was assessed using a two-sided, two-sample *t*-test with permutation-based FDR < 0.001 and $S0$ = 0.1 (Dataset S1).

**(e)** Category enrichment analysis of nucleolar-enriched proteins from (d). Enriched categories were identified using Fisher's exact test with a Benjamini–Hochberg FDR cut-off of < 0.001, enrichment factor > 2, and intersection size > 2 (Dataset S2). Each circle represents an enriched category from Gene Ontology Cellular Component (GOCC) or *Dorner et al.* ribosome biogenesis annotations[1]; circle size indicates the number of intersecting proteins.

**(f)** Pie chart of nucleolar-enriched proteins identified by ANOVA (Dataset S3), showing the proportion of proteins whose nucleolar enrichment was regulated in a RAS–MAPK-dependent manner, a RAS-dependent but MAPK-independent manner, or was unaffected by $Kras^{G12D}$ expression. Numbers of proteins in each category are indicated in brackets.

**(g)** Category enrichment analysis of proteins recruited to the nucleolus upon $Kras^{G12D}$ induction, in a MAPK-independent manner, from (Figure 2b). Enriched categories were identified using Fisher's exact test with a Benjamini–Hochberg FDR cut-off of < 0.001, enrichment factor > 2, and intersection size > 2 (Dataset S5). Each circle represents an enriched category from Gene Ontology Cellular Component (GOCC) or *Dorner et al.* ribosome biogenesis annotations[1]; circle size indicates the number of intersecting proteins.

**(h)** Category enrichment analysis of proteins lost from the nucleolus upon $Kras^{G12D}$ induction, in a MAPK-dependent manner, from (Figure 2b). Enriched categories were identified using Fisher's exact test with a Benjamini–Hochberg FDR cut-off of < 0.001, enrichment factor > 2, and intersection size > 2 (Dataset S5). Each circle represents an enriched category from Gene Ontology Cellular Component (GOCC) or *Dorner et al.* ribosome biogenesis annotations[1]; circle size indicates the number of intersecting proteins.

**(i)** Category enrichment analysis of proteins lost from the nucleolus upon $Kras^{G12D}$ induction, in a MAPK-independent manner, from (Figure 2b). Enriched categories were identified using Fisher's exact test with a Benjamini–Hochberg FDR cut-off of < 0.001, enrichment factor > 2, and intersection size > 2 (Dataset S5). Each circle represents an enriched category from Gene Ontology Cellular Component (GOCC) or *Dorner et al.* ribosome biogenesis annotations[1]; circle size indicates the number of intersecting proteins.

**(j)** PCA of phosphoproteomic profiles obtained from nucleolar fractions and their matched WCLs of iKras cells, under the indicated conditions. Samples show a clear separation between nucleolar and WCL, as well as an overall clustering of biological replicates within different treatment group. The red arrow indicates an outlier WCL sample which was removed from subsequent analysis.

**(k)** Volcano plot comparison of the combined nucleolar fractions versus WCL phospho-samples from (j). Significance was assessed using a two-sided, two-sample t-test, using an FDR cut off of 0.001 and an S0 of 0.1 (Dataset S6).

**(l)** Category enrichment analysis of nucleolar-enriched phosphorylated proteins from (k). Enriched categories were identified using Fisher's exact test with a Benjamini–Hochberg FDR cut-off of < 0.001, enrichment factor > 2, and intersection size > 2 (Dataset S7). Each circle represents an enriched category from Gene Ontology Cellular Component (GOCC) or *Dorner et al.* ribosome biogenesis annotations[1]; circle size indicates the number of intersecting proteins.

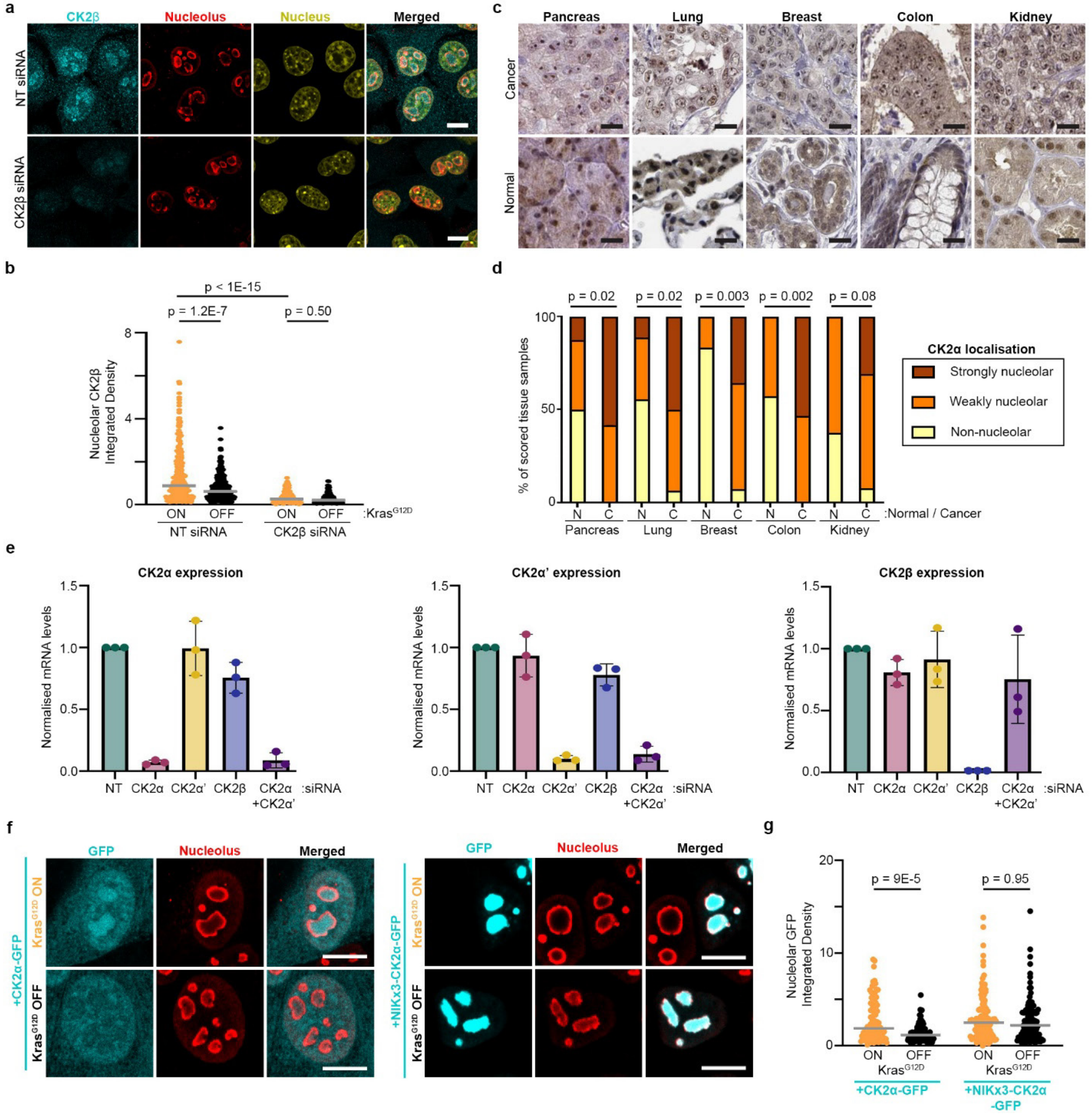


**Supplementary Data Figure S3: CK2 localises to the nucleolus across diverse cancers.**

**(a)** Validation of CK2β antibody specificity for immunofluorescence analysis. Representative immunofluorescence images of CK2β (cyan) in iKras cells treated with non-targeting control (NT) or CK2β-targeting siRNAs for 72 h, with doxycycline added for the last 24 h to induce Kras$^{G12D}$ expression. Nucleoli were visualised by Ncl immunostaining (red). DNA was stained with Hoechst (yellow). Scale bar, 10 µm.

**(b)** Quantification of nucleolar CK2β levels from the immunofluorescence images shown in (a). Quantifications were from iKras cells treated with non-targeting control (NT) or CK2β-targeting siRNAs for 72 h, with or without doxycycline addition during the final 24 h to induce Kras$^{G12D}$ expression. A total of n =

993 nucleoli pooled from three independent biological replicates were analysed. Statistical significance was assessed by one-way ANOVA with Dunn's multiple comparisons test.

**(c)** Representative CK2α immunohistochemistry (IHC) images from normal and malignant pancreas, lung, breast, colon, and kidney tissues, extracted from Human Protein Atlas (https://www.proteinatlas.org/), showing nucleolar enrichment of CK2α in malignant tissues. Scale bar, 20 μm.

**(d)** Quantification of nucleolar CK2α localisation in the IHC images from (c). Images were scored as 'non-nucleolar' if no specific signal enrichment in the nucleolus could not be seen, 'weekly nucleolar' if nucleolar staining could be seen in a minority of the cells, and strongly nucleolar if nucleolar staining could be seen in most of the cells. 8 to 10 IHC slides, each from an independent patient, were scored.

**(e)** Confirmation of subunit-specific CK2 knockdowns, alone or in combination, by specific pools of siRNAs. iKras cells were transfected with the indicated pools of siRNAs for 72 h, before lysis, RNA extraction, and RT-qPCR analysis with specific probes against CK2α (left), CK2α' (middle), or CK2β (right). CK2 subunit mRNA levels were normalised to Gapdh as loading control.

**(f)** Representative immunofluorescence images of iKras cells ectopically expressing GFP-tagged wild-type CK2α (left) or NIKx3-fused CK2α (right), with or without doxycycline treatment for 24 h to induce $Kras^{G12D}$ expression. GFP was used to visualise the ectopic CK2α proteins (cyan). Nucleoli were visualised by Ncl immunostaining (red). Nuclei were stained with Hoechst (yellow). Scale bar, 10 μm.

**(g)** Quantification of nucleolar GFP-tagged CK2α levels from the immunofluorescence images shown in (f). NIKx3-fusion successfully traps CK2 in the nucleolus irrespective of $Kras^{G12D}$ expression status. A total of n = 402 nucleoli pooled from three independent biological replicates were analysed. Significance was assessed by one-way ANOVA with Dunn's multiple comparisons testing.

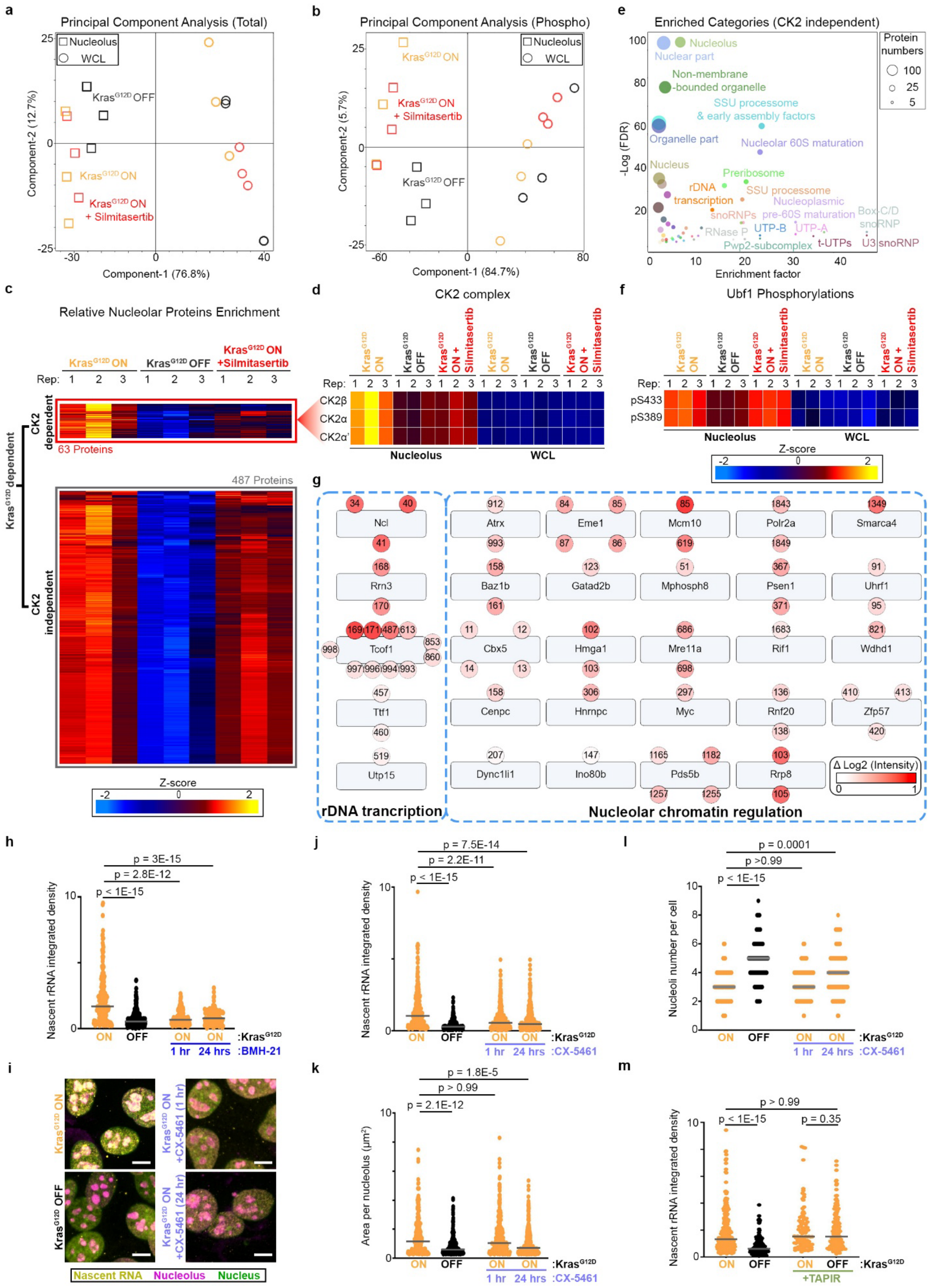
a
Principal Component Analysis (Total)
Nucleolus
WCL
Kras^G12D OFF
Kras^G12D ON
Kras^G12D ON + Silmitasertib
Component-1 (76.8%)
Component-2 (12.7%)
b
Principal Component Analysis (Phospho)
Component-1 (84.7%)
Component-2 (5.7%)
e
Enriched Categories (CK2 independent)
Protein numbers
Nucleolus
Nuclear part
Non-membrane -bounded organelle
SSU processome & early assembly factors
Organelle part
Nucleolar 60S maturation
Nucleus
Preribosome
rDNA transcription
SSU processome
Nucleoplasmic pre-60S maturation
snoRNPs
Box C/D snoRNP
RNase P
UTP-B
UTP-A
Pwp2-subcomplex
t-UTPs
U3 snoRNP
Enrichment factor
-Log (FDR)
c
Relative Nucleolar Proteins Enrichment
Kras^G12D ON
Kras^G12D OFF
Kras^G12D ON +Silmitasertib
Rep:
CK2 dependent
63 Proteins
487 Proteins
CK2 independent
Kras^G12D dependent
Z-score
d
CK2 complex
CK2β
CK2α
CK2α'
Nucleolus
WCL
f
Ubf1 Phosphorylations
pS433
pS389
Z-score
g
Ncl
Rrn3
Tcof1
Ttf1
Utp15
rDNA trancription
Atrx
Eme1
Mcm10
Polr2a
Smarca4
Baz1b
Gatad2b
Mphosph8
Psen1
Uhrf1
Cbx5
Hmga1
Mre11a
Rif1
Wdhd1
Cenpc
Hnrnpc
Myc
Rnf20
Zfp57
Dync1li1
Ino80b
Pds5b
Rrp8
Δ Log2 (Intensity)
Nucleolar chromatin regulation
h
p = 3E-15
p = 2.8E-12
p < 1E-15
Nascent rRNA integrated density
ON OFF ON ON :Kras^G12D
1 hr 24 hrs :BMH-21
i
Kras^G12D ON
Kras^G12D OFF
Kras^G12D ON +CX-5461 (1 hr)
Kras^G12D ON +CX-5461 (24 hr)
Nascent RNA Nucleolus Nucleus
j
p = 7.5E-14
p = 2.2E-11
p < 1E-15
Nascent rRNA integrated density
ON OFF ON ON :Kras^G12D
1 hr 24 hrs :CX-5461
k
p = 1.8E-5
p > 0.99
p = 2.1E-12
Area per nucleolus (μm²)
ON OFF ON ON :Kras^G12D
1 hr 24 hrs :CX-5461
l
p = 0.0001
p >0.99
p < 1E-15
Nucleoli number per cell
ON OFF ON ON :Kras^G12D
1 hr 24 hrs :CX-5461
m
p > 0.99
p < 1E-15
p = 0.35
Nascent rRNA integrated density
ON OFF ON OFF :Kras^G12D
+TAPIR

**Supplementary Data Figure S4: Nucleolar CK2 phosphorylates rDNA chromatin and RNAPI transcription machinery to boost rRNA synthesis and nucleolar fusion.**

**(a)** PCA of total proteomic profiles obtained from nucleolar fractions and their matched WCLs of iKras cells, under the indicated conditions.

**(b)** PCA of phosphoproteomic profiles obtained from nucleolar fractions and their matched WCLs of iKras cells, under the indicated conditions.

**(c)** Heatmap of proteins whose nucleolar levels relative to the WCL was significantly increased upon 24 h of $Kras^{G12D}$ induction in iKras cells, in a CK2-dependent or CK2-independent manner, as judged by sensitivity to 1 h of Silmitasertib (10 μM) treatment. Significant changes were identified using one-way ANOVA with permutation-based FDR of < 0.05 and S0 = 0.1, followed by post hoc testing with FDR of < 0.05 (Dataset S10).

**(d)** Heatmap showing relative levels of CK2α, CK2α, and CK2β, in the nucleolus and WCL samples of indicated treatment conditions. Localisation of CK2 subunits to the nucleolus is sensitive to Silmitasertib.

**(e)** Category enrichment analysis of $Kras^{G12D}$-dependent CK2-independent nucleolar phosphorylation targets from (Figure 4a). Enriched categories were identified using Fisher's exact test with a Benjamini–Hochberg FDR cut-off of < 0.02, enrichment factor > 2, and intersection size > 2 (Dataset S14). Each circle represents an enriched category from Gene Ontology Cellular Component (GOCC) or *Dorner et al.* ribosome biogenesis annotations[1]; circle size indicates the number of intersecting proteins.

**(f)** Heatmap showing relative levels of Ubf1 S433 and S389 phosphorylations in the nucleolus and WCL samples of indicated treatment conditions. Ubf1 phosphorylations are sensitive to $Kras^{G12D}$, but not Silmitasertib treatment.

**(g)** Selected $Kras^{G12D}$ and CK2-dependent nucleolar phosphorylations belonging to rDNA transcription and nucleolar chromatin-related protein categories. Colour difference highlights the degree of change in log2 intensity between $Kras^{G12D}$ and $Kras^{G12D}$ + Silmitasertib treatment conditions.

**(h)** Quantification of nucleolar FUrd levels from the immunofluorescence images shown in (Figure 4j). A total of n = 907 nucleoli pooled from three independent biological replicates were analysed. Significance was assessed by one-way ANOVA with Dunn's multiple comparisons testing.

**(i)** Representative nascent RNA immunofluorescence images of iKras cells treated with or without doxycycline for 24 h to induce $Kras^{G12D}$ expression, with or without CX-5461 (0.1 μM) co-treatment for indicated times to inhibit rRNA synthesis. Cells were pulse-labelled with FUrd (2 mM) for 30 min to label nascent RNA, before immunostaining for FUrd (yellow). Nucleoli were visualised by Ncl immunostaining (red). Nuclei were stained with Hoechst (green). Scale bar, 10 μm.

**(j)** Quantification of nucleolar FUrd levels from the immunofluorescence images shown in (i). A total of n = 1370 nucleoli pooled from three independent biological replicates were analysed. Significance was assessed by one-way ANOVA with Dunn's multiple comparisons testing.

**(k)** Quantification of individual nucleolar area from the immunofluorescence images shown in (i). A total of n = 1,370 nucleoli pooled from three independent biological replicates were analysed. Significance was assessed by one-way ANOVA with Dunn's multiple comparisons testing.

**(l)** Quantification of nucleolar number per cell from the immunofluorescence images shown in (i). A total of n = 416 cells pooled from three independent biological replicates were analysed. Significance was assessed by one-way ANOVA with Dunn's multiple comparisons testing.

**(m)** Quantification of nucleolar FUrd levels from the immunofluorescence images shown in (Figure 4n). A total of n = 701 nucleoli pooled from three independent biological replicates were analysed. Significance was assessed by one-way ANOVA with Dunn's multiple comparisons testing.

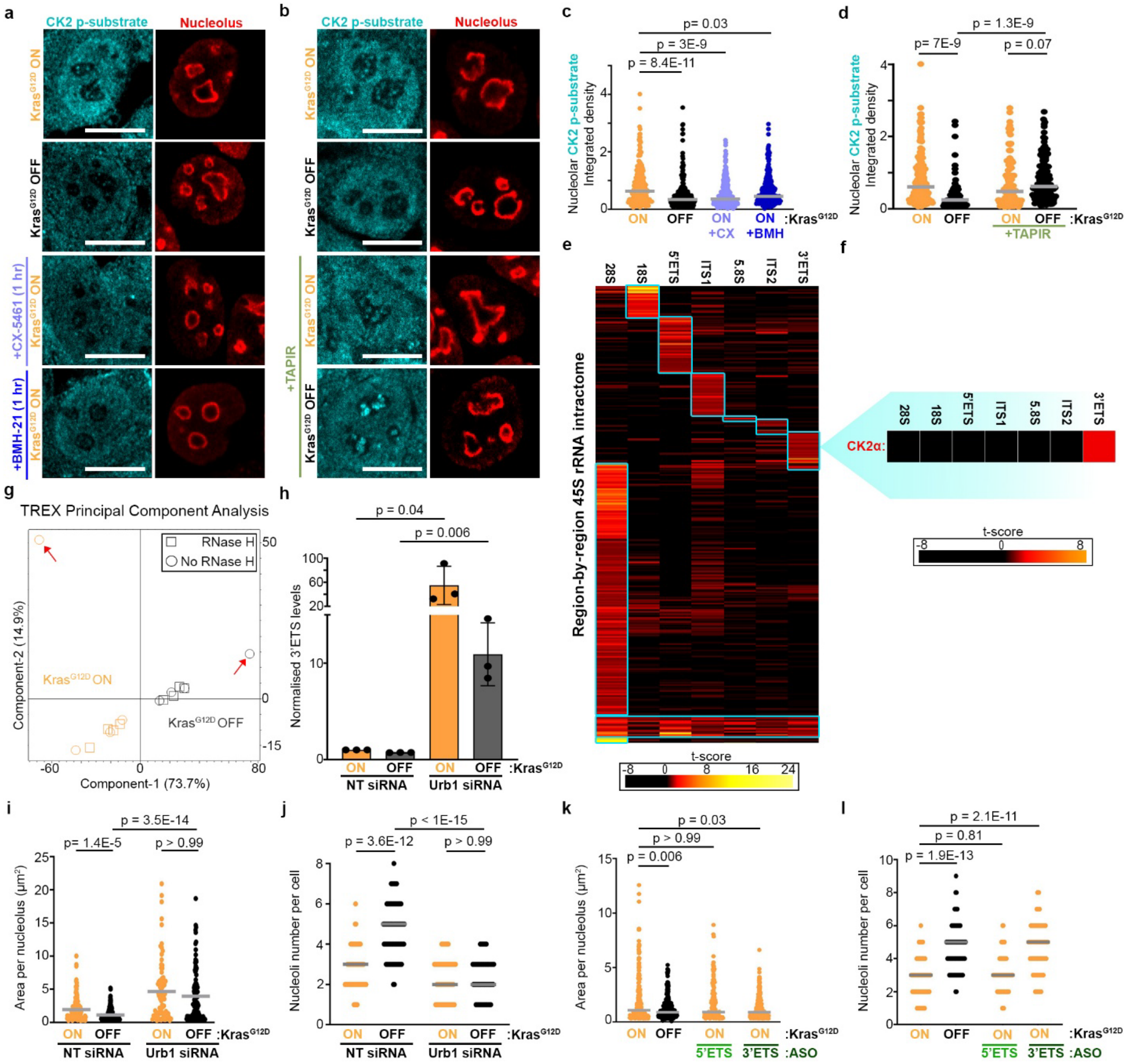


**Supplementary Data Figure S5: 3'ETS rRNA localises CK2 signalling to the nucleolus.**

**(a)** Representative immunofluorescence images of CK2 phosphorylated substrates (cyan) in iKras cells treated with or without doxycycline for 24 h to induce Kras$^{G12D}$ expression, or with doxycycline plus 1 h co-treatment with CX-5461 (100 nM) or BMH-21 (500 nM) to inhibit RNAPI. Nucleoli were visualised by Ncl immunostaining (red). Scale bar, 10 μm.

**(b)** Representative immunofluorescence images of CK2 phosphorylated substrates (cyan) in iKras cells treated with or without doxycycline for 24 h to induce Kras$^{G12D}$ expression, with or without TAPIR expression to ectopically activate RNAPI. Nucleoli were visualised by Ncl immunostaining (red). Scale bar, 10 μm.

**(c)** Quantification of nucleolar levels of CK2 phosphorylated substrates from the immunofluorescence images shown in (a). A total of n = 1,104 nucleoli pooled from three independent biological replicates were analysed. Significance was assessed by one-way ANOVA with Dunn's multiple comparisons testing.

**(d)** Quantification of nucleolar levels of CK2 phosphorylated substrates from the immunofluorescence images shown in (b). A total of n = 543 nucleoli pooled from three independent biological replicates were analysed. Significance was assessed by one-way ANOVA with Dunn's multiple comparisons testing.

**(e)** Clustering analysis of TREX-identified protein interactors of different human 45S rRNA regions, from Dodel et al.[2], revealing clusters of proteins enriched as specific interactors with the indicated rRNA regions. Clustering with complete Euclidean distance and K-means preprocessing was performed on the TREX t-scores from +RNase H vs. No RNase H control t-test analyses for each region.

**(f)** Zoomed-in view of CK2α from (e), showing binding specifically to the 3'ETS region.

**(g)** PCA of 3'ETS TREX samples (+RNase H and No RNase H control) from iKras cells grown with or without doxycycline for 24 h to induce $Kras^{G12D}$ expression. The red arrow indicates outlier no RNase H control samples, which were removed from further downstream analysis.

**(h)** RT-qPCR analysis of 3′ETS RNA levels in iKras cells transfected with Urb1-targeting or non-targeting control siRNAs, with or without doxycycline for 24 h to induce $Kras^{G12D}$ expression. Cells were subjected to lysis, RNA extraction, and RT-qPCR analysis with 3′ETS-specific probes. The 3'ETS levels were normalised to β-Actin mRNA as a house-keeping loading control. Significance was assessed by two-tailed unpaired t-test (n = 3 independent biological replicates).

**(i)** Quantification of individual nucleolar area from the immunofluorescence images shown in (Figure 5j). A total of n = 504 nucleoli pooled from three independent biological replicates were analysed. Significance was assessed by one-way ANOVA with Dunn's multiple comparisons testing.

**(j)** Quantification of nucleolar number per cell from the immunofluorescence images shown in (Figure 5j). A total of n = 261 cells pooled from three independent biological replicates were analysed. Significance was assessed by one-way ANOVA with Dunn's multiple comparisons testing.

**(k)** Quantification of individual nucleolar area from the immunofluorescence images shown in (Figure 5l). A total of n = 985 nucleoli pooled from three independent biological replicates were analysed. Significance was assessed by one-way ANOVA with Dunn's multiple comparisons testing.

**(l)** Quantification of nucleolar number per cell from the immunofluorescence images shown in (Figure 5l). A total of n = 270 cells pooled from three independent biological replicates were analysed. Significance was assessed by one-way ANOVA with Dunn's multiple comparisons testing.

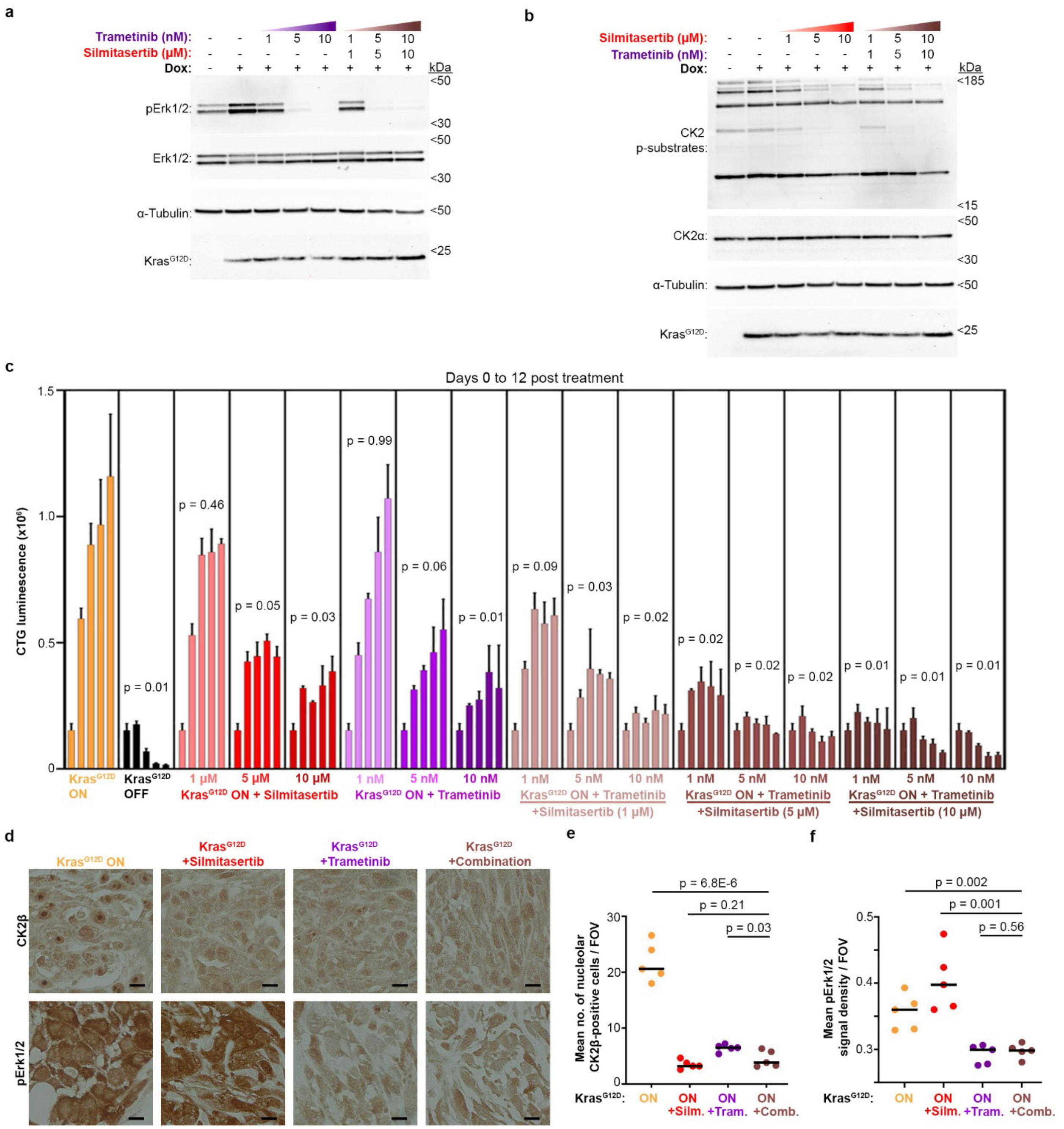


**Supplementary Data Figure S6: CK2 and MAPK inhibitor combination blocks *in vitro* anchorage-independent PDAC cell proliferation and *in vivo* tumour growth.**

**(a)** Western blot confirmation of RAS–MAPK signalling inhibition in iKras cells by Trametinib treatment, alone or in combination with Silmitasertib, at indicated drug concentrations. iKras cells were grown with or without doxycycline for 24 h to induce $Kras^{G12D}$ expression, or with doxycycline along with co-treatment with the indicated inhibitor doses for 1 h, before lysis and analysis by western blotting with the indicated antibodies.

**(b)** Western blot confirmation of CK2 signalling inhibition in iKras cells by Silmitasertib treatment, alone or in combination with Trametinib, at indicated drug concentrations. iKras cells were grown with or without doxycycline for 24 h to induce $Kras^{G12D}$ expression, or with doxycycline along with co-treatment with the indicated inhibitor doses for 1 h, before lysis and analysis by western blotting with the indicated antibodies.

**(c)** Quantification of spheroid viability from (Figure 6e). Spheroids were established for 3 days from 2,000 iKras cells seeded per well of 96-well low attachment plates in the presence of doxycycline. They were subsequently subjected to the indicated treatments for up to 12 days. Viability was measured at 0, 3, 6, 9, and 12 days post-establishment, using the 3D CellTiter-Glo (CTG) reagent assay. CTG luminescence values were plotted as measures of cell viability. A total of n = 4 independently seeded spheroids per timepoint per condition were analysed. Statistical significance was assessed by two-way ANOVA with Dunnett's multiple comparisons test. Reported p-values correspond to the 12-day spheroids.

**(d)** Representative IHC images of endpoint xenograft PDAC tumours from (Figure 6g), stained for CK2β (top) or pErk1/2 (bottom). Scale bar, 10 μm.

**(e)** Quantification of nucleolar CK2β staining from images shown in (d). Average number of nucleolar CK2β positive cells per field of view (FOV) for each individual tumour was quantified. A total of 5 averaged FOVs from n = 5 independent tumours per condition were quantified. Significance was assessed using a two-tailed unpaired t-test.

**(f)** Quantification of overall pErk1/2 staining from images shown in (d). Average pErk1/2 optical density per field of view (FOV) for each individual tumour was quantified. A total of 5 averaged FOVs from n = 5 independent tumours per condition were quantified. Significance was assessed using a two-tailed unpaired t-test.

### Supplementary Dataset Legends:

**Dataset S1:** Source data for the t-test comparison of nucleolar vs. WCL proteomic samples, from Supplementary Data Fig. S2d. Significance was calculated using a two-sided two-sample t-test analysis, with permutation-based FDR cutoff of < 0.001 and S0 = 0.1. Proteins significantly enriched in the nucleolus are marked in the "*Significant? (nucleolar)*" column.

**Dataset S2:** Source data for the category enrichment analysis of nucleolar proteins, from Supplementary Data Fig. S2e. Significance was calculated using Fisher's exact test analysis with a Benjamini–Hochberg FDR cut-off of < 0.001, enrichment factor > 2, and intersection size > 2.

**Dataset S3:** Source data for ANOVA coupled with post-hoc comparison of nucleolar and WCL proteomic samples with or without $Kras^{G12D}$ induction, or with $Kras^{G12D}$ induction and Trametinib co-treatment. Significance was calculated using a one-way ANOVA analysis with a permutation-based FDR cutoff of < 0.05 and S0 = 0.1, followed by post-hoc Tukey HSD comparison (FDR <0.05). Proteins significantly enriched in the nucleolus are marked in the "*nucleolar (posthoc)*" column. Significant post-hoc comparisons are listed in the "*Posthoc Significant pairs (0.05 FDR)*" column.

**Dataset S4:** Source data for ANOVA coupled with post-hoc comparison of $Kras^{G12D}$-regulated nucleolar proteins, from Fig. 2b. Post-hoc clusters from Dataset S3 corresponding to significantly increasing or decreasing nucleolar proteins in a $Kras^{G12D}$ and MAPK-dependent manner, or $Kras^{G12D}$-dependent but MAPK-independent manner, were selected.

**Dataset S5:** Source data for category enrichment analysis of Kras$^{G12D}$-sensitive nucleolar proteins, from Fig. 2c and Supplementary Fig. S2g–i. Significance was calculated using Fisher's exact test analysis with a Benjamini–Hochberg FDR cut-off of < 0.001, enrichment factor > 2, and intersection size > 2.

**Dataset S6:** Source data for the t-test comparison of nucleolar vs. WCL phosphoproteomic samples, from Supplementary Data Fig. S2k. Significance was calculated using a two-sided two-sample t-test analysis, with permutation-based FDR cutoff of < 0.001 and S0 = 0.1. Phospho-sites significantly enriched in the nucleolus are marked in the "*Significant? (nucleolar)*" column.

**Dataset S7:** Source data for the category enrichment analysis of nucleolar phospho-proteins, from Supplementary Data Fig. S2l. Significance was calculated using Fisher's exact test analysis with a Benjamini–Hochberg FDR cut-off of < 0.001, enrichment factor > 2, and intersection size > 2.

**Dataset S8:** Source data for the normalised enrichment scores of serine/threonine kinase motifs that change in the nucleolus upon Kras$^{G12D}$ induction, from Fig. 2e. Motif enrichment analysis was performed using the Kinase Library motif enrichment tool, using an FDR cutoff of < 0.05 (https://kinase-library.phosphosite.org/kinase-library/motif-enrichment-analysis)[3].

**Dataset S9:** Source data for the normalised enrichment scores of serine/threonine kinase motifs that change in the WCL upon Kras$^{G12D}$ induction, from Fig. 2f. Motif enrichment analysis was performed using the Kinase Library motif enrichment tool, using an FDR cutoff of < 0.05 (https://kinase-library.phosphosite.org/kinase-library/motif-enrichment-analysis)[3].

**Dataset S10:** Source data for ANOVA coupled with post-hoc comparison of nucleolar and WCL proteomic samples, with or without Kras$^{G12D}$ induction, or with Kras$^{G12D}$ induction and short-term (1 h) Silmitasertib co-treatment. Significance was calculated using a one-way ANOVA analysis with a permutation-based FDR cutoff of < 0.05 and S0 = 0.1, followed by post-hoc Tukey HSD comparison (FDR <0.05). Significant post-hoc comparisons are listed in the "*Posthoc Significant pairs (0.05 FDR)*" column.

**Dataset S11:** Source data for ANOVA coupled with post-hoc comparison of nucleolar and WCL phosphoproteomic samples, with or without Kras$^{G12D}$ induction, or with Kras$^{G12D}$ induction and short-term (1 h) Silmitasertib co-treatment. Significance was calculated using a one-way ANOVA analysis with a permutation-based FDR cutoff of < 0.05 and S0 = 0.1, followed by post-hoc Tukey HSD comparison (FDR <0.05). Phospho-sites significantly enriched in the nucleolus are marked in the "*Nucleolar? (posthoc)*" column. Significant post-hoc comparisons are listed in the "*Posthoc Significant pairs (FDR<0.05)*" column.

**Dataset S12:** Source data for ANOVA coupled with post-hoc comparison of Kras$^{G12D}$-regulated nucleolar phospho-sites, from Fig. 4a. Post-hoc clusters from Dataset S11 corresponding to significantly increasing nucleolar phospho-sites in a Kras$^{G12D}$ and CK2-dependent manner, or Kras$^{G12D}$-dependent but CK2-independent manner, were selected.

**Dataset S13:** Source data for the serine/threonine kinase motif enrichment in the Kras$^{G12D}$ and CK2-dependent nucleolar phospho-sites, from Fig. 4b. Motif enrichment was performed using the Kinase Library Fisher Enrichment Analysis tool, with an FDR cutoff of < 0.05 (https://kinase-library.phosphosite.org/kinase-library/fisher-enrichment-analysis)[3].

**Dataset S14:** Source data for category enrichment analysis of Kras$^{G12D}$ and CK2-dependent nucleolar phospho-proteins, from Fig. 4c. Significance was calculated using Fisher's exact test analysis with a Benjamini–Hochberg FDR cut-off of < 0.02, enrichment factor > 2, and intersection size > 2.

**Dataset S15:** Source data of the list of identified proteins in the *3'ETS* TREX experiments, with or without Kras$^{G12D}$ expression, from Fig. 5g and 5h. Significance was calculated using two-sided two-sample t-tests, with a p–value cutoff of < 0.02 and an absolute difference score > 0.5. Significant hits in each condition are marked by '+'
